# Selective vulnerability of cerebral vasculature to *NOTCH3* variants in small vessel disease and rescue by phosphodiesterase-5 inhibitor

**DOI:** 10.1101/2025.05.30.656914

**Authors:** Xiangjun Zhao, Chaowen Yu, Antony Adamson, Aite Zhao, Huiyu Zhou, Pankaj Sharma, Tao Wang

## Abstract

*NOTCH3* variants cause CADASIL the most common genetic form of small vessel disease (SVD) and vascular dementia (VaD) and increase the stroke and SVD/VaD risk. CADASIL is a systemic vasculopathy but predominantly manifests in the brain. The molecular mechanisms of CADASIL remain largely unclear with no specific available treatments. *NOTCH3* is primarily expressed in vascular smooth muscle cells (VSMCs). Using human induced pluripotent stem cell (iPSC) models and developmental lineage-specific VSMC differentiation, we revealed a selective vulnerability of cerebral but not peripheral VSMC mimics to *NOTCH3* variants. Transcriptomic, protein and functional analyses demonstrated a switching of CADASIL iPSC-VSMCs from a contractile to a synthetic phenotype, accompanied with extensive extracellular matrix accumulation and impairment of cell adhesion leading to *anoikis*. Importantly, we describe an endothelial independent nitric oxide signalling in VSMCs which was dysregulated in the CADASIL iPSC-VSMCs, and posphodiesterase-5 (PDE5) inhibition successfully rescued the abnormal VSMC function, suggesting a novel therapeutic strategy. Our findings offer new mechanistic insights into brain specific phenotype in *NOTCH3*-associated SVD/VaD and support patient-specific iPSCs to be a valuable model for identifying targeted treatment for *NOTCH3*-associated SVD/VaD.

## Introduction

Small vessel disease (SVD) is the leading cause of vascular dementia (VaD) and the second commonest cause of dementia after Alzheimer’s disease^1^. Genetic factors play an important role in the susceptibility, development and progression of SVD^2,3^. *NOTCH3* variants cause CADASIL (cerebral autosomal dominant arteriopathy and subcortical infarcts and leukoencephalopathy), the most common form of genetic SVD^4,5^. This rare genetic form of SVD has an estimated prevalence of 2-4 per 100,000 individuals, but CADASIL-like *NOTCH3* variants are more common than expected with an overall frequency of ∼1:300 in the general population and associated with a significantly increased risk of sporadic stroke and SVD/VaD^6–8^. CADASIL is a systemic vasculopathy affecting small vessels throughout the body, but its clinical manifestations are mainly in the brain. Patients with CADASIL typically experience recurrent strokes, and many also suffer from migraine with aura and mood disturbance gradually progressing to cognitive impairment and eventually, vascular dementia (VaD)^9–11^. The exact molecular mechanisms by which *NOTCH3* variants lead to neurological symptoms remain unclear; and consequently, no specific or effective treatments are currently available.

*NOTCH3* is one of the four Notch receptors transducing Notch signalling predominantly expressed in arterial vascular smooth muscle cells (VSMCs), where it promotes differentiation and proliferation, prevents migration, and protects VSMCs against apoptosis^12–16^, playing a key role in maintaining vascular homeostasis. Unlike Notch1 or Notch2, depletion of which is embryonically lethal, *Notch3* knockout mice are viable but display abnormal VSMC maturation and venous characteristics^13^. In CADASIL, the hallmark pathological changes are VSMC degeneration, NOTCH3 protein accumulation, and granular osmophilic material (GOM) deposition in small arteries^10,16–18^. However, *Notch3* knockout mice do not have typical CADASIL phenotype^13^, and RBP-Jκ mediated canonical Notch signalling remains active in transgenic CADASIL mice^19^, suggesting the disease is not due to a simple loss of function. Instead, toxic gain-of-function from protein aggregation likely drives disease progression^9^. Given that Notch signalling cross-talks with multiple signalling pathways, including the Wnt^20,21^, hedgehog (Hg)^22^, transforming growth factor-beta (TGF-β)^23^, PI3K/Akt^21,24^, MAPK/ERK^25^, and nitric oxide (NO)^26^ signalling pathways, it remains unclear whether and in what way mutant *NOTCH3* disrupts the crosstalk and contributes to the development of CADASIL pathology. Understanding this mechanism could have important therapeutic implications.

VSMCs are developmentally heterogeneous^27,28^. Neural crest cells form VSMCs in the ascending aorta and cerebral vessels; lateral plate mesoderm contributes to VSMCs in the aortic root and coronary arteries via epicardium; paraxial mesoderm generates VSMCs in peripheral vessels. Lineage-specific VSMCs have differential responses to growth factors like angiotensin II^29^ and TGF-β^30^, and show varying disease susceptibilities, independent of hemodynamic factors; for instance, Marfan’s syndrome affects the aorta root and ascending aorta^31,32^, while atherosclerotic targets the aortic arch and coronary arteries^33,34^. *NOTCH3* is systemically expressed throughout in vasculature, its lineage-specific susceptibility to *NOTCH3* variants remain unexplored, leaving a key knowledge gap in understanding the brain-predominant clinical features in CADASIL.

VSMCs are not terminally differentiated cells and exhibit phenotypic plasticity in response to different stimuli^35,36^. In healthy arteries they exhibit a contractile phenotype, but under pathological conditions like atherosclerosis, vascular injury or restenosis, they undergo dedifferentiation and acquire a synthetic phenotype by downregulating contractile markers and increasing production of extracellular matrix (ECM) proteins, growth factors and cytokines, leading to proliferation and migration^27,28^. In CADASIL, VSMC phenotype has not been systemically studied, despite impaired vasoreactivity observed in both patients and transgenic mice^37–39^. A deeper understanding of VSMC plasticity in CADASIL is crucial to uncover disease mechanisms and informing therapeutic strategies.

To date, over a dozen CADASIL mouse models have been developed, which uncovered valuable insights into molecular mechanisms of the disease. While the models successfully phenocopy key pathological features of CADASIL, such as VSMC degeneration, GOM deposition and Notch3 accumulation, most failed to recapitulate the full range of neurological symptoms observed in human patients^40^. Given that 90% of drugs that pass preclinical testing in animal models ultimately fail to reach patients^41^, there is a critical need for human disease models as complimentary tools for studying disease mechanisms and advancing drug development. In this regard, induced pluripotent stem cells (iPSCs) derived from CADASIL patients provide a powerful platform for better understanding of this condition and accelerating therapeutic discovery.

We previously established iPSC models of CADASIL from patient skin biopsies, revealing that CADASIL VSMCs failed to stabilise angiogenic capillary structures and support blood-brain barrier (BBB) integrity^42,43^. In this study, we generated developmental lineage-specific VSMCs and identified a selective vulnerability of neural crest derived VSMCs to *NOTCH3* variants, which likely explain the brain-specific manifestation in CADASIL. Results provide convincing evidence of VSMC phenotype switching and demonstrate accumulations of ECM proteins and impaired VSMC adhesion leading to *anoikis* type of cell death. The local NO signalling in CADASIL was impaired with reduced soluble guanylyl cyclase (sGC) in the CADASIL iPSC-VSMCs. Treatment with an NO donor or phosphodiesterase type 5 (PDE5) inhibitor (sildenafil) significantly reversed VSMC dysfunction and cell death, suggesting a promising therapeutic approach for CADASIL.

## Results

### IPSC-derived brain-specific VSMCs are selectively vulnerable to *NOTCH3* variants in CADASIL

To determine if VSMCs in the brain vasculature have specific susceptibility to the *NOTCH3* variants in CADASIL which change cell behaviour and functions, we differentiated iPSCs into VSMCs via neural crest (NC-SMCs), lateral plate mesoderm (LPM-SMCs), and paraxial mesoderm (PM-SMCs), which represent VSMCs of the brain, the heart, and peripheral, respectively, using established methods^32,44^ with modifications (**Figs. 1a**).

**Figure 1.**
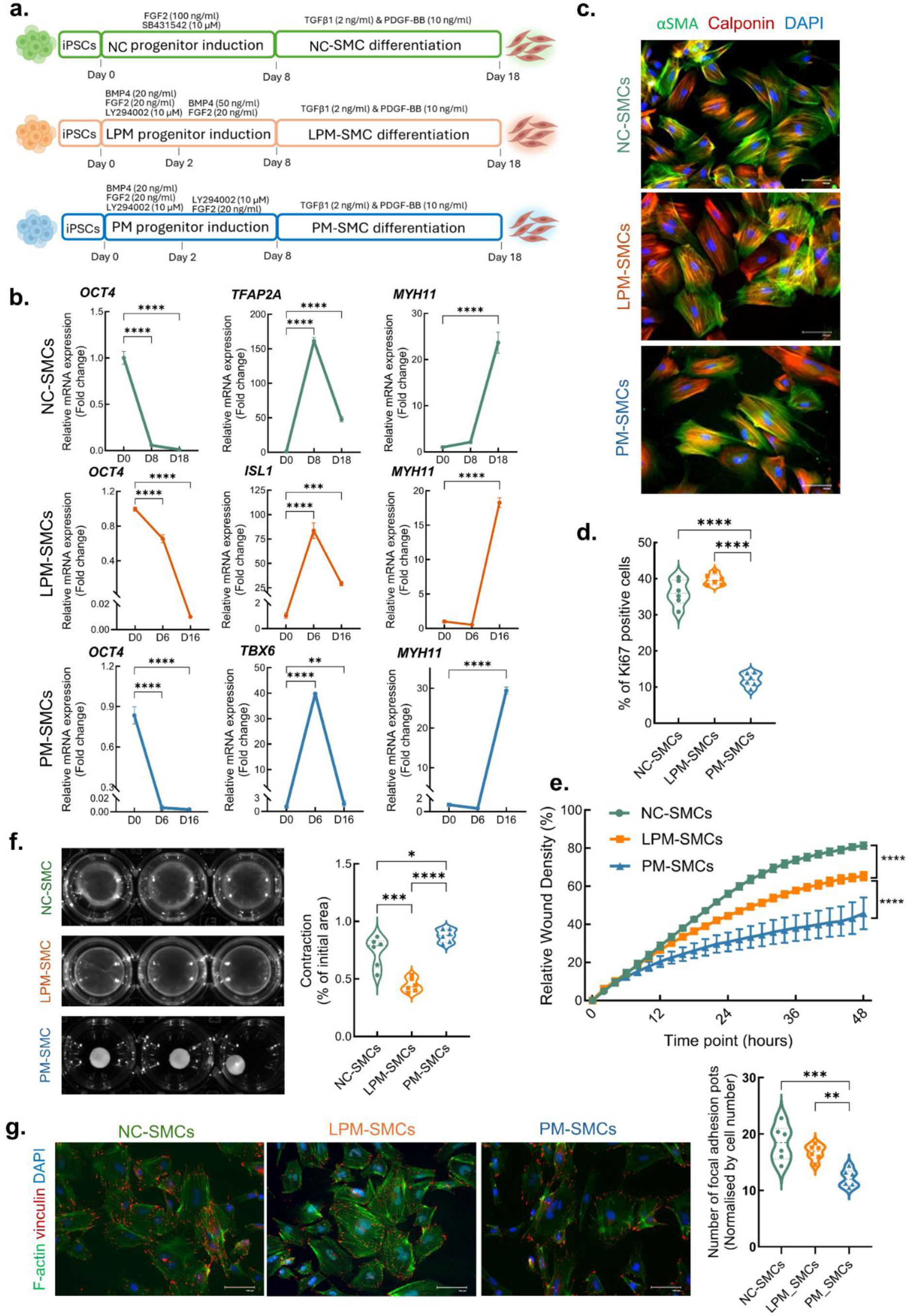
Lineage-specific VSMC differentiation from iPSCs. **a** Diagram illustrating protocols used for differentiating iPSCs into VSMCs via neural crest lineage (NC-SMCs), lateral plate mesoderm lineage (LPM-SMCs) and paraxial mesoderm lineage (PM-SMCs). **b** RT-qPCR determination of marker gene expressions showing downregulation of pluripotency marker gene *OCT4* during VSMC differentiation, upregulation of the lineage-specific maker genes, *TFAP2A*, *ISL1* and *TBX6,* at the respective progenitor stages, and increased expression of VSMC marker gene *MYH11* in differentiated VSMCs. **c** IPSC-derived lineage-specific VSMCs on differentiation day 18 were immunofluorescent stained with VSMC markers, α-SMA (green) and calponin (red), and nuclear counterstained by DAPI (blue). **d** Proliferation of lineage-specific VSMCs was determined by ki67 immunofluorescent staining. **e** Migration of lineage-specific VSMCs was determined by IncuCyte live cell imaging of wound healing assay. **f** Contraction of lineage-specific VSMCs was determined by collagen I gel contraction assay and quantified (right panel). **g** Vinculin immunofluorescent staining (red) of iPSC derived VSMCs showing focal adhesions and its quantitation (right panel). F-actin was stained green. Scale bars,100μm in **c** and **g**. Data are presented as mean ± SEM from at least 3 independent iPSC differentiations (n=3). One-way ANOVA and Tukey’s post hoc test in **b**, **d**, **f** and **g,** or two-way ANOVA before post hoc test demonstrating overall differences between groups in **e**, *p ≤ 0.05, **p ≤ 0.01, ***p ≤ 0.001, ****p ≤ 0.0001.

We first used a wild-type iPSC line to validate features of the origin-specific iPSC-SMCs. During the course of differentiation, pluripotency markers were downregulated, and developmental lineage-specific markers *TFAPA2* (neural crest), *ISL1* (lateral plate mesoderm), and *TBX6* (paraxial mesoderm) were significantly upregulated at the corresponding progenitor stages (**Fig 1b**). The three sub-types of iPSC-SMCs all expressed VSMC specific markers including *ACTA2*/α-SMA, *CNN1*/calponin, *MYH11*/SMMHC and *TAGLN*/SM22α as determined by RT-qPCR (**Fig. 1b**) and immunofluorescent staining (**Fig. 1c and S1a**). Meanwhile, each type of the VSMCs expressed the corresponding lineage-specific markers: *SEMA3α* and *MEF2c* for NC-SMCs, *ISL1* and *HAND1* for LPM-SMCs, and *HOXA10* and *HOXC6* for PM-SMCs (**Fig. S1b**)^45–47^. Comparing to PM-SMCs, the NC-SMCs and LPM-SMCs had much higher proliferation and migration rates, with the NC-SMCs being the highest, as measured by Ki67 immunofluorescence staining (**Fig. 1d and S2a**) and wound healing assay (**Fig. 1e and S2b**), respectively. In contrast, PM-SMCs exhibited the highest contractility, and the LPM-SMCs were least contractile as measured by collagen I contraction assay (**Fig. 1f**). These behavioural and functional properties of the origin-specific iPSC-VSMCs align well with findings reported on primary VSMCs obtained from various vascular locations^48,49^. Additionally, we found for the first time that the NC-SMCs exhibited the highest number of focal adhesions, followed by the LPM-SMCs, as measured by vinculin staining (**Fig. 1g**), suggesting a more active interaction between VSMCs and the matrix in the vasculature of the CNS. PM-SMCs displayed the least number of focal adhesions (**Fig. 1g**).

Having validated the identity of the lineage-specific iPSC-VSMCs, we compared the behaviour and basic functionalities between VSMCs derived from iPSCs of two CADASIL patients (R153C and C224Y) and their corresponding isogenic controls (isoCtrls) that we have made recently using CRISPR/Cas9 gene editing (**Fig. S3**). Cell proliferation as determined by Ki67 staining revealed a significant increase in NC-SMCs derived from the two CADASIL iPSC lines as compared to their isogenic controls (**Fig. 2a**). Interestingly, neither LPM-SMCs nor PM-SMCs from the two CADASIL iPSC lines had significantly altered their proliferation rate, suggesting a specific change in brain VSMCs in CADASIL. Cell migration was conducted using the IncyCyte wound healing assay, revealing again that the CADASIL NC-SMCs, but not the LPM-SMCs nor PM-SMCs, had significantly increased migration compared to the isoCtrls (**Fig. 2b and S4**), further suggesting a selective susceptibility of CNS VSMCs to the CADASIL *NOTCH3* variants.

**Figure 2.**
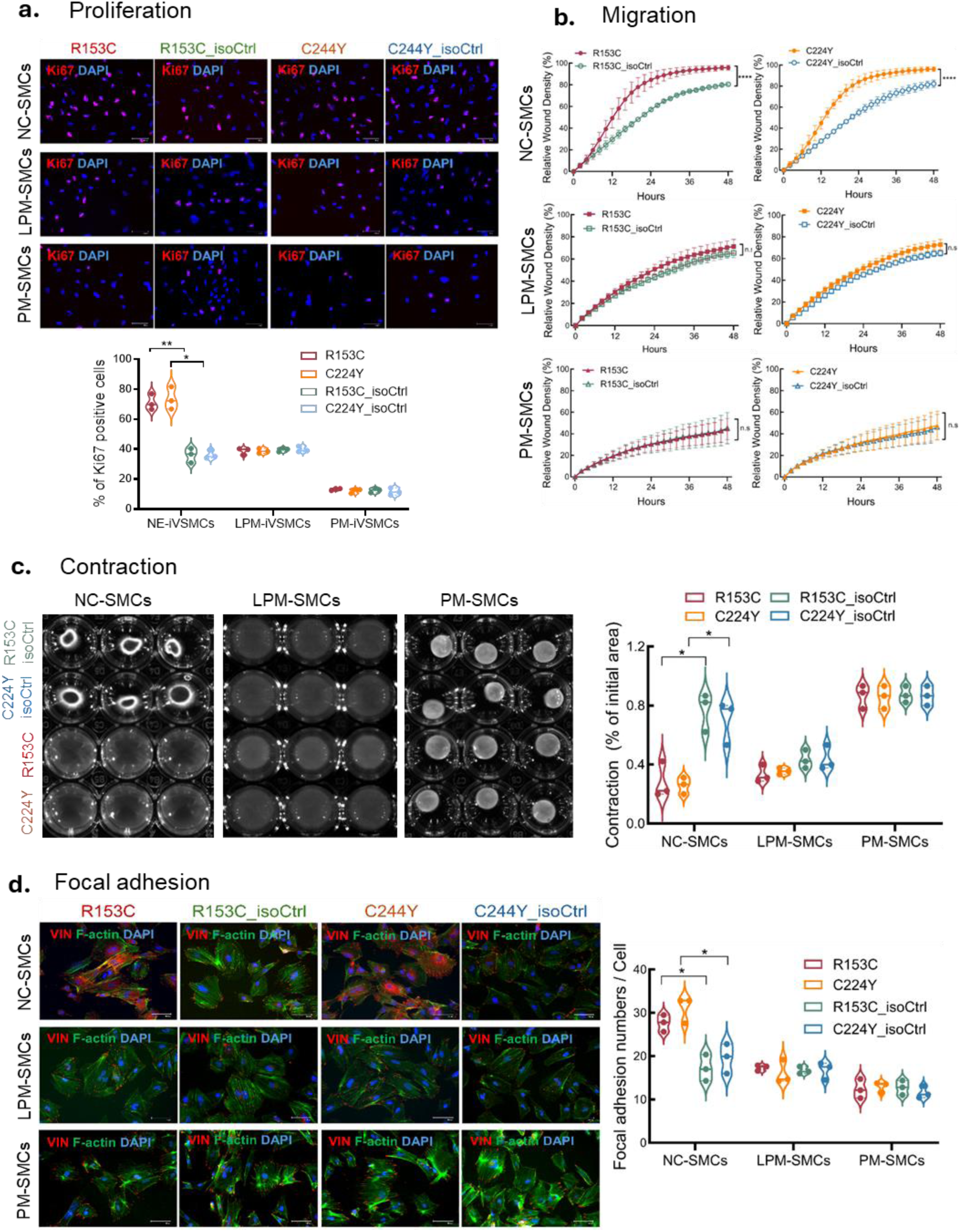
Comparison of behaviours and functions between lineage-specific VSMCs derived from CADASIL iPSCs. IPSCs from two CADASIL patients (R153C & C224Y) and their isogenic controls (isoCtrl) were differentiated into lineage-specific VSMCs: NC-SMCs, LPM-SMCs and PM-SMCs, respectively. **a** Cell proliferation was determined by immunofluorescent staining of Ki67 (red). Nuclei were counter stained by DAPI (blue). Results were quantified and showed in the lower panel. Scale bars = 100 μm. **b** Cell migration was determined by IncuCyte live cell imaging of wound healing assay. **c** Cell contraction was determined by collagen I contraction assay which was quantified (right panel). **d** Focal adhesion was visualised by vinculin immunofluorescent staining (red) which was quantified (right panel). F-actin was stained green. Scale bars = 100 μm in **a** and **d**. All quantifiable data are presented as mean ± SEM from 3 independent iPSC differentiations (n=3). Unpaired Student’s *t* test in **a**, **c** and **d**. Two-way ANOVA before post-hoc test demonstrating overall differences between groups in **b**, *p ≤ 0.05, **p ≤ 0.01, ****p ≤ 0.0001.

In addition to the behavioural changes, CADASIL NC-SMCs exhibited significantly impaired contractile function in the collagen I gel contraction assay, unlike the LPS-SMCs and PM-SMCs that showed no difference from their isogenic controls (**Fig. 2c**). Vinculin immunostaining revealed increased but disorganised focal adhesions in CADASIL NC-SMCs, along with diffused cytosolic vinculin signals (**Fig. 2d**), suggesting potential defects in vinculin activation and membrane recruitment^50,51^, likely disrupting cytoskeletal organisation^52^.

Taken together, results from comparing the behaviour and function of the developmental lineage-specific VSMCs derived from iPSCs strongly suggest a brain-specific vascular vulnerability to the CADASIL *NOTCH3* variants (**Fig. 2**). In the following experiments, the iPSC NC-SMCs model, namely iVSMCs, was exclusively used to study disease mechanisms.

### CADASIL iVSMCs undergo phenotype switching from contractile to synthetic state

To gain a more comprehensive understanding of genes and signalling pathways that are involved in the pathological changes in CADASIL and identify novel disease mechanisms and therapeutic targets, we conducted bulk RNA sequencing (RNAseq) on NC-SMCs differentiated from two CADASIL iPSC lines (R153C and C224Y), two isogenic control lines (R153C_isoCtrl and C224Y_isoCtrl), and three wild-type control lines (02C9, 02C3 and EIPL1). Principal component analysis (PCA) showed that PC1 explained 67% of the variability that mainly distinguished the CADASIL samples from all the controls (**Fig. 3a**). Differential expression analysis revealed 4161 differentially expressed genes (p-value < 0.05 & Log ^(Fold^ ^change)^ > 1), including 1932 upregulated and 2229 downregulated genes, between the CADASIL and control samples (**Fig. 3b**). As the isoCtrls and wild-type controls clustered well together on the PCA plot (**Fig. 3a**), thus, only the isogenic controls were used in the subsequent data analysis in order to more accurately identify the impact by the *NOTCH3* variants and avoid the influence by inter-person variations (**Fig. 3c**). Gene ontology (GO) analysis on GO terms of biological process (BP), cellular components (CC) and molecular function (MF) highlighted changes including extracellular matrix organisation, focal adhesion, smooth muscle cell proliferation, smooth cell migration, cell-matrix adhesion, actin filament organisation, collagen-containing extracellular matrix, and regulation of vasculature development (**Fig. 3d**), suggesting a matrix related pathology and phenotype change of CADASIL VSMCs.

**Figure 3.**
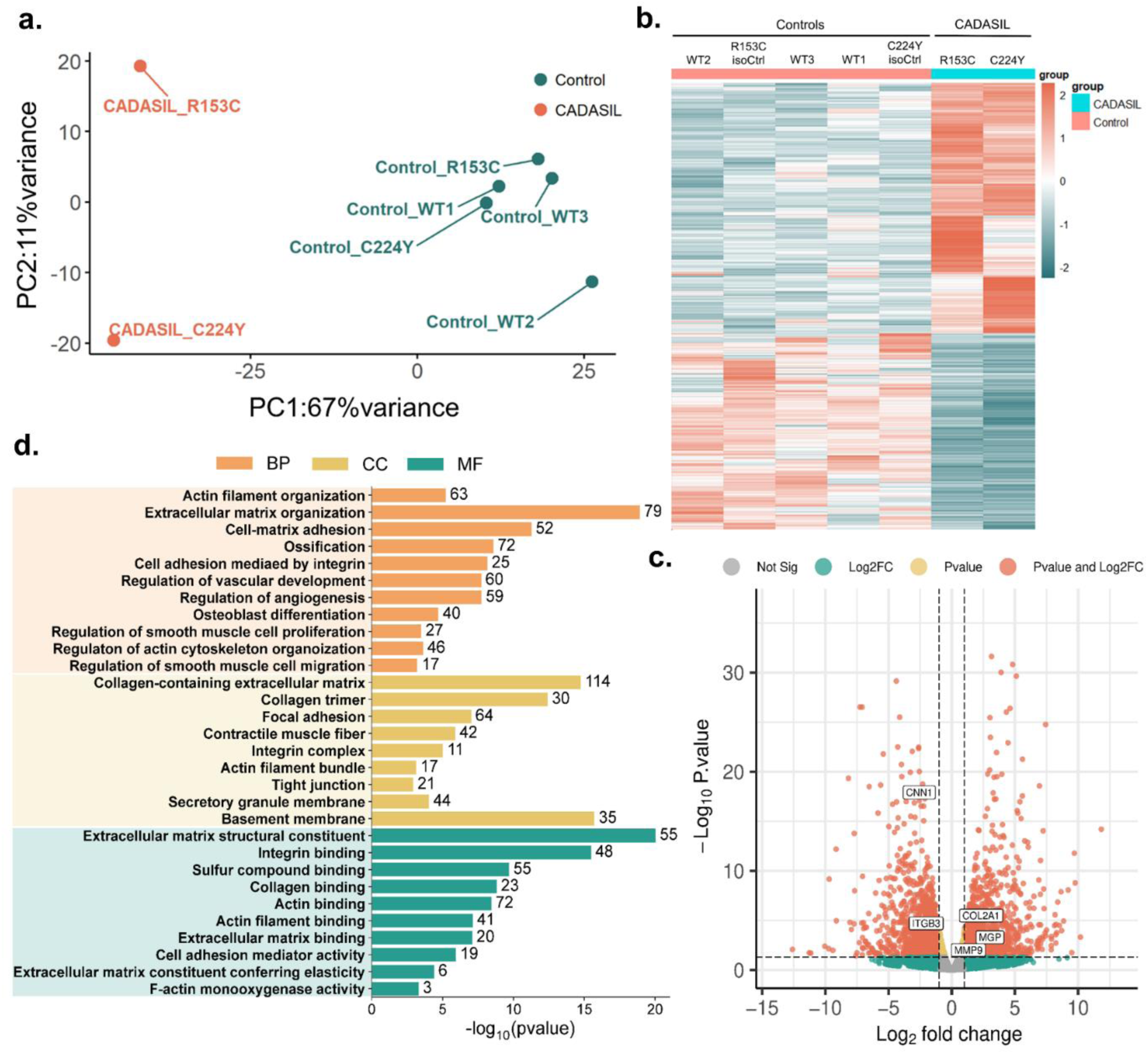
Analyses of RNA sequencing data. IPSCs from two CADASIL patients (R135C & C224Y) and respective isogenic controls (isoCtrl) as well as three wild-type iPSC control lines (WT1-3) were differentiated into iVSMCs via neural crest lineage (NC) which were subjected to RNA sequencing. **a** Principal component (PCA) plot. **b** Heatmap showing differentially expressed genes (DEGs) between iVSMCs of the two patients and five controls (2x isoCtrls and 3x WT controls). **c** Volcano plot comparing DEGs between CADASIL patients (R135C & C224Y) and their isogenic controls (R153C_isoCtrl and C224Y_isoCtrl). **d** GO (gene ontology) term enrichment analysis of DEGs from the RNAseq data. BP, biological processes, CC, cellular components, MF, molecular functions.

We then specifically analysed VSMC phenotype (**Fig. 4a**) on the iPSC CADASIL model. Heatmap of the RNAseq data shows downregulation of a range of contractile marker genes including *MYH11*, *ACTA2*, *CNN1*, *TAGLN,* and *MYOCD*, and upregulation of many synthetic marker genes including *COL1A2*, *MMP9*, *SPP1*, *MYH10*, and *MSN* in the CADASIL iVSMCs (**Fig. 4b**) comparing to isoCtrls, which was confirmed by RT-qPCR (**Fig. 4c**). The downregulation of key contractile markers, α-SMA, MLCK, calponin, and SM22-α, in CADASIL iVSMCs were also confirmed by western blotting (**Fig. 4d**). A chord diagram shows downregulation of a range of genes that linked to GO terms of actin filament binding and muscle cell contraction (**Fig. 4e**). Increased proliferation and migration and reduced contractility are functional features of synthetic VSMCs^36^, which we have demonstrated on CADASIL iVSMCs above in **Fig. 2**. Additional migration assay was conducted using 3D VSMC spheroids, further confirming the increased migration of CADASIL iVSMCs (**Fig. S5)**. Taken together, results from gene expressions, protein levels and functional assays suggest that CADASIL iVSMCs undergo phenotype switching from a contractile to a synthetic state.

**Figure 4.**
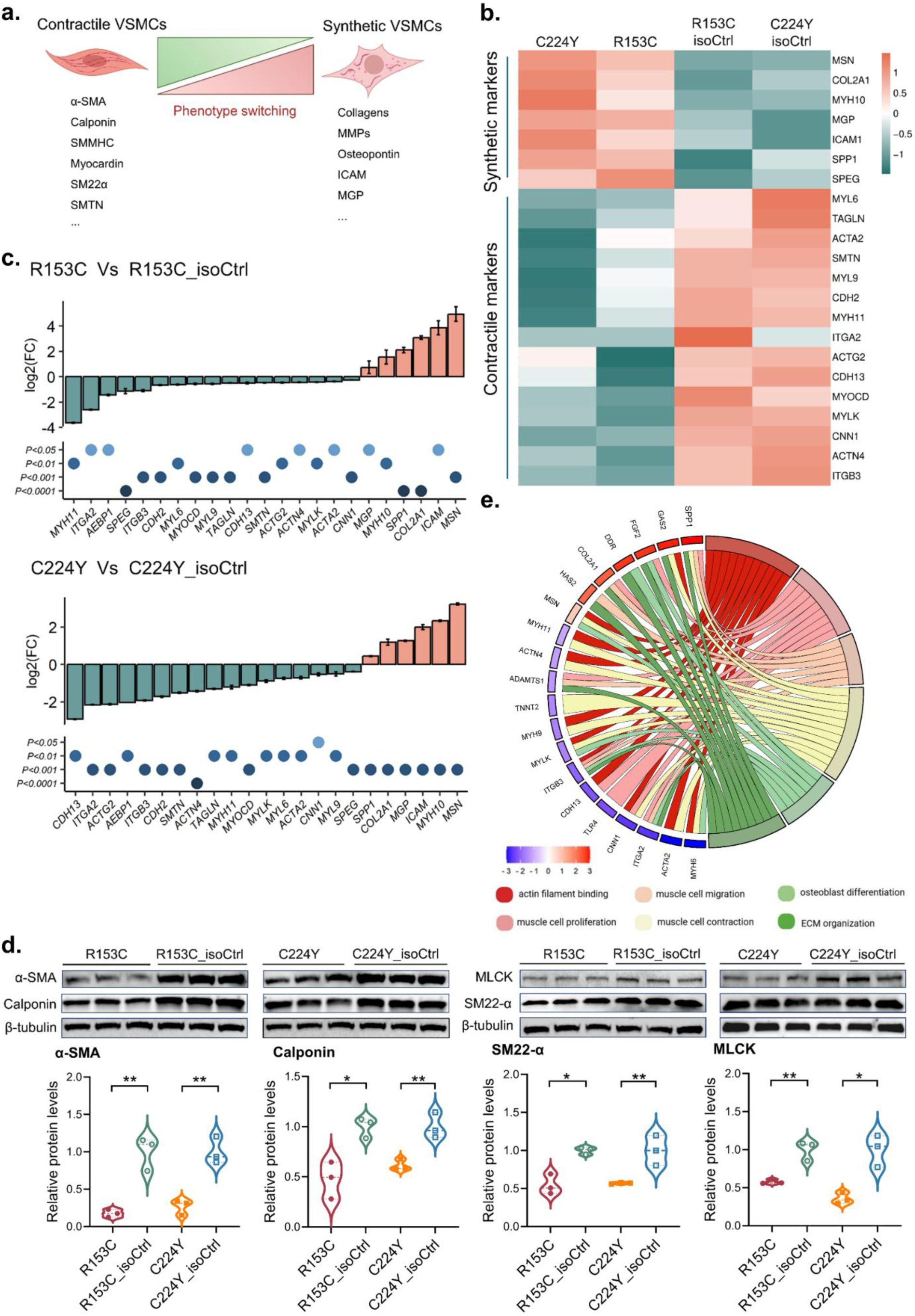
Phenotype switching of iPSC derived CADASIL iVSMCs. IPSCs from two CADASIL patients (R135C & C224Y) and their isogenic controls (R153C_isoCtrl and C224Y_isoCtrl) were differentiated into iVSMCs via neural crest (NC). **a** Diagram illustrating principles of VSMC switching from a contractile to a synthetic phenotype and associated key marker gene expressions. **b** Heatmap showing differentially expressed VSMC contractile and synthetic marker genes from RNA sequencing (RNAseq) results of the two CADASIL and their isogenic control iVSMCs. **c** RT-qPCR confirmation of the up- and down-regulations of major VSMC contractile and synthetic marker genes in CADASIL iVSMCs. Data are presented as mean ± SEM from 3 independent iPSC differentiations (n=3). Fold change of gene expressions for each gene were calculated separately and compared by unpaired Student’s *t* test. **d** Western blotting (WB) confirmation of downregulations of key VSMC contractile proteins, α-SMA, calponin, SM22α and MLCK, in CADASIL iVSMCs, comparing to their isogenic controls. Quantifications of the WB results show in the panel below. Data are presented as mean ± SEM from 3 independent iPSC differentiations with 3 technical replicants in each experiment (n=3). Unpaired Student’s *t* test, *p ≤ 0.05, **p ≤ 0.01. **e** Chord diagram generated showing relationship between some of the VSMC phenotype related Gene Ontology (GO) terms and differentially expressed genes (DEGs). Diagram **a** was Created with BioRender.com.

### ECM composition and adhesion-actin coupling are severely disrupted in CADASIL iVSMCs leading to cell death

Under the gene ontology (GO) term “extracellular matrix organisation (p = 3.49E-24)”, there were around 150 differentially expressed between the CADASIL iPSC-NC-SMCs and isogenic controls as shown on the heatmap (**Fig. S6a**). Out of the 150 genes, 43 genes were identified to be collagen related, with the majority of which were upregulated (**Fig. S6b**). We then conducted Gene Set Enrichment Analysis (GSEA) to uncover the coordinated change of genes involved in ECM function. GSEA revealed that gene sets relating to “ECM structural constituent”, “collagen biosynthesis and modifying enzymes”, and “collagen trimer” were positively enriched in the CADASIL iVSMCs (**Fig. 5a**), suggesting matrix accumulation and its abnormal turnover in CADASIL. RT-qPCR results confirmed the significant upregulation of key basement membrane components including collagen subtypes (*COL4A2, COL1A1, COL1A2, COL2A1* and *COL3A1*), MMP (*MMP9*), TIMPs (*TIMP1* and *TIMP3*), and *FN* (fibronectin) in the CADASIL iVSMCs comparing to isoCtrls (**Fig. 5b**). Interestingly, MMP3 was consistently found downregulated in the CADASIL iVSMCs (**Fig. 5b**). Considering the unique role of MMP3 that has much broader ECM substrates, including fibronectin and proteoglycans apart from collagens, than other MMPs and participates in proMMP activations^53^, the reduced MMP3 may play a significant role in the ECM remodelling in CADASIL. Consistent with the transcriptomic and RT-qPCR results, spheroids formed from CADASIL iVSMCs displayed collagen IV accumulation (**Fig. 5c**). TEM images revealed large amount of matrix deposition (**Fig. 5d**), which is consistent with findings on autopsy samples of human patient ^54^. Accompanied with the matrix deposition, accumulation of Notch3 extracellular domain proteins, the hallmark of CADASIL pathology, was also faithfully recapitulated on the iVSMC spheroids (**Fig. 5e**) and western blotting (**Fig. 5f**). Despite accumulation, the mutant Notch3 had impaired CBF1/RBPJκ-mediated canonical Notch activity as determined by luciferase assay (**Fig. S7**).

**Figure 5.**
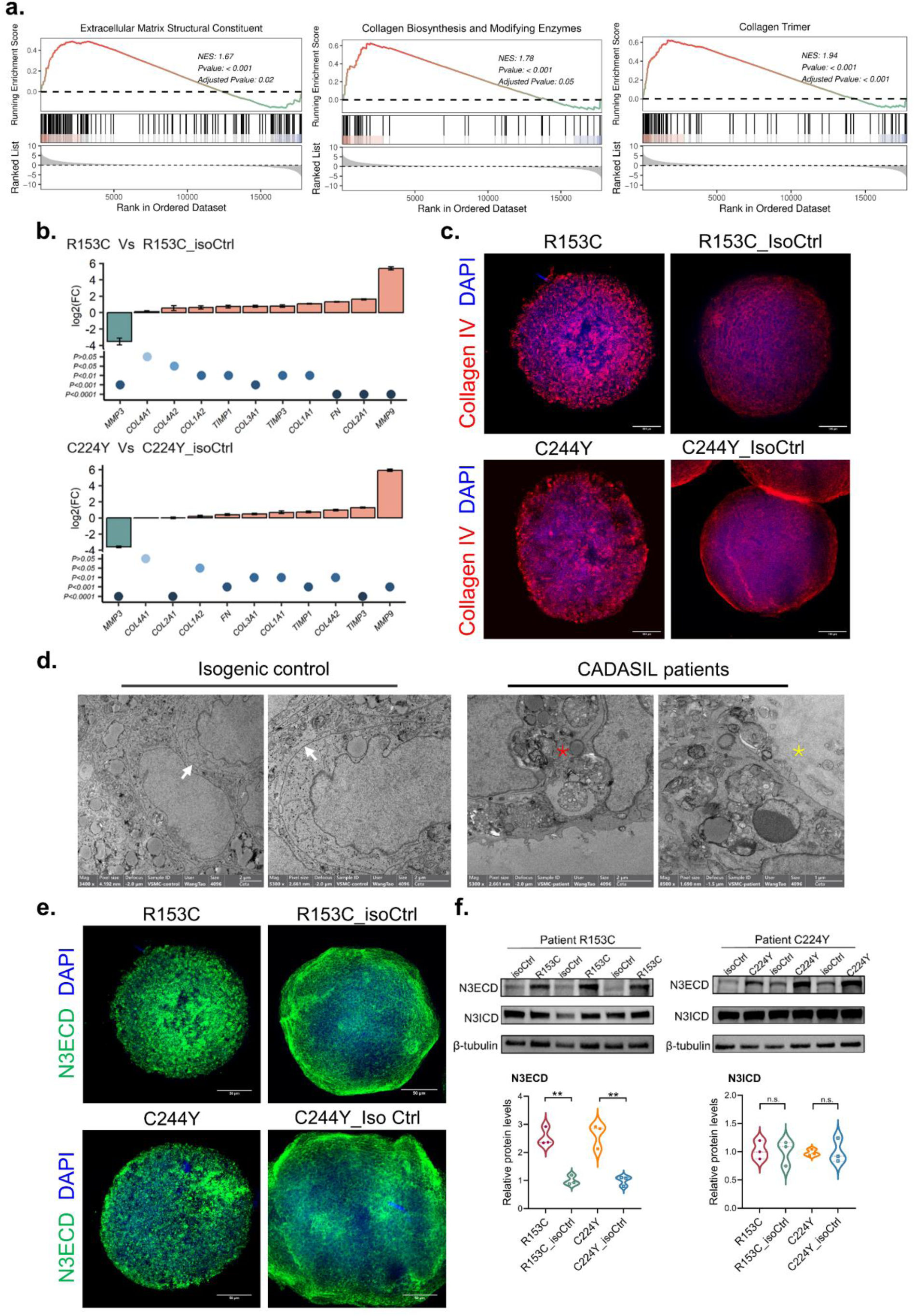
Extracellular matrix (ECM) accumulation in CADASIL iVSMCs. IPSCs from two CADASIL patients (R135C & C224Y) and their isogenic controls (R153C_isoCtrl and C224Y_isoCtrl) were differentiated into iVSMCs via neural crest (NC) lineage. **a** Gene set enrichment analysis (GSEA) plot of RNA sequencing data on gene ontology (GO) categories of ECM structural constituent, collagen biosynthesis and modifying enzymes, and collagen trimer in CADASIL iVSMCs versus their isogenic controls. **b** RT-qPCR confirmation of differentially expressed major extracellular matrix genes in CADASIL iVSMCs versus isogenic controls. Data are presented as mean ± SEM from 3 independent iPSC differentiations (n=3). Fold changes of gene expression for each gene were calculated separately and compared by unpaired Student’s *t* test. **c** Collagen IV immunofluorescent staining of iVSMCs spheroids derived from iPSCs of CADASIL and isoCtrls. Nuclei were counterstained by DAPI. Scale bar = 50μm. **d** TEM images of iVSMCs derived from iPSCs of CADASIL and isoCtrls. White arrows, smooth cell membranes between adjacent cells. Red asterisk star, vanish of cell junctions and accumulation of extracellular materials in CADASIL iVSMCs. Yellow asterisk star, ECM accumulation. **e** Notch3 extracellular domain (N3ECD) immunofluorescent staining of iVSMCs spheroids derived from iPSCs of CADASIL and isoCtrls. Nuclei were counterstained by DAPI. Scale bar = 50μm. **f** Western blotting (WB) of iVSMCs using antibodies against N3ECD or Notch3 intracellular domain (N3ICD). Quantifications of the WB results are shown in the lower panel. Data are presented as mean ± SEM from 3 independent iPSC differentiations (n=3). Unpaired Student’s *t* test, **p ≤ 0.01.

KEGG analysis highlighted pathways relating to “cytoskeleton in muscle cells”, “focal adhesion”, “regulation of cytoskeleton”, “cell adhesion molecules”, and “ECM-receptor interaction” (**Fig. 6a**), suggesting changes in focal adhesion-actin coupling in CADASIL iVSMCs. Key components of focal adhesion include integrins, tailin, vinculin, paxillin, and focal adhesion kinase (FAK), which link actin cytoskeleton to ECM via integrins (**Fig. 6b**), playing a crucial role in cell-matrix interaction regulating cell behaviours and survival. We then quantified the expression of a range of integrin subunits (*ITGA1, ITGA2, ITGA3, ITGA4, ITGA5, ITGA6, ITGA7, ITGB1, ITGB3*) in the iVSMCs using RT-qPCR and found all of which were significantly downregulated in the CADASIL iPSC-NC-SMCs, except for *ITGA4* that was upregulated in the mutant cells (**Fig. 6c**). Western blotting demonstrated significant reduction in total FAK and pFAK as well as pFAK/FAK ratio in the CADASIL iVSMCs (**Fig. 6d**), suggesting a likely disruption of cell-matrix interaction which compromises cell survival^55^. Fluorescent staining of F-actin cytoskeleton revealed disorganised actin fibre (**Fig. 6e**). To quantify this abnormality, we developed a computer vision algorithm based on Canny edge detection and measures of the disorder of samples, which demonstrated a significantly altered spatial dispersion of actin filaments in the CADASIL iVSMCs as compared to the isoCtrls (**Fig. 6f**), implicating compromised contractility, which is in line with the damaged contractile function (**Fig. 2c**) and reduced contractile markers (**Fig. 4**) in CADASIL iVSMCs as described earlier. Additionally, GSEA showed significantly negative enrichment of “adherence junction” gene sets (**Fig. 6g**), suggesting additional defect in cell-cell interactions. This is also revealed on the TEM images, where smooth lines of cell membranes were observed between adjacent isoCtrl iVSMCs, which was hardly seen in the CADASIL iVSMCs (**Fig. 5d**). Instead, large amounts of cell debris and matrix were frequently observed between adjacent CADASIL iVSMCs (**Fig. 5d**). Taken together, our evidence on the disruption of interactions between both cell-to-matrix and cell-to-cell suggests a loss of cell anchorage, likely leading to *anoikis*^56^. Indeed, live/dead assay demonstrated a significant increase of cell death in the CADASIL iVSMCs (**Figs. 6h and 6i**). *IGTA4* has specific role in leukocyte adhesion and ECM remodelling in a form of α4β1^57^, the upregulation of which (**Fig. 6c**) warrants further investigation.

**Figure 6.**
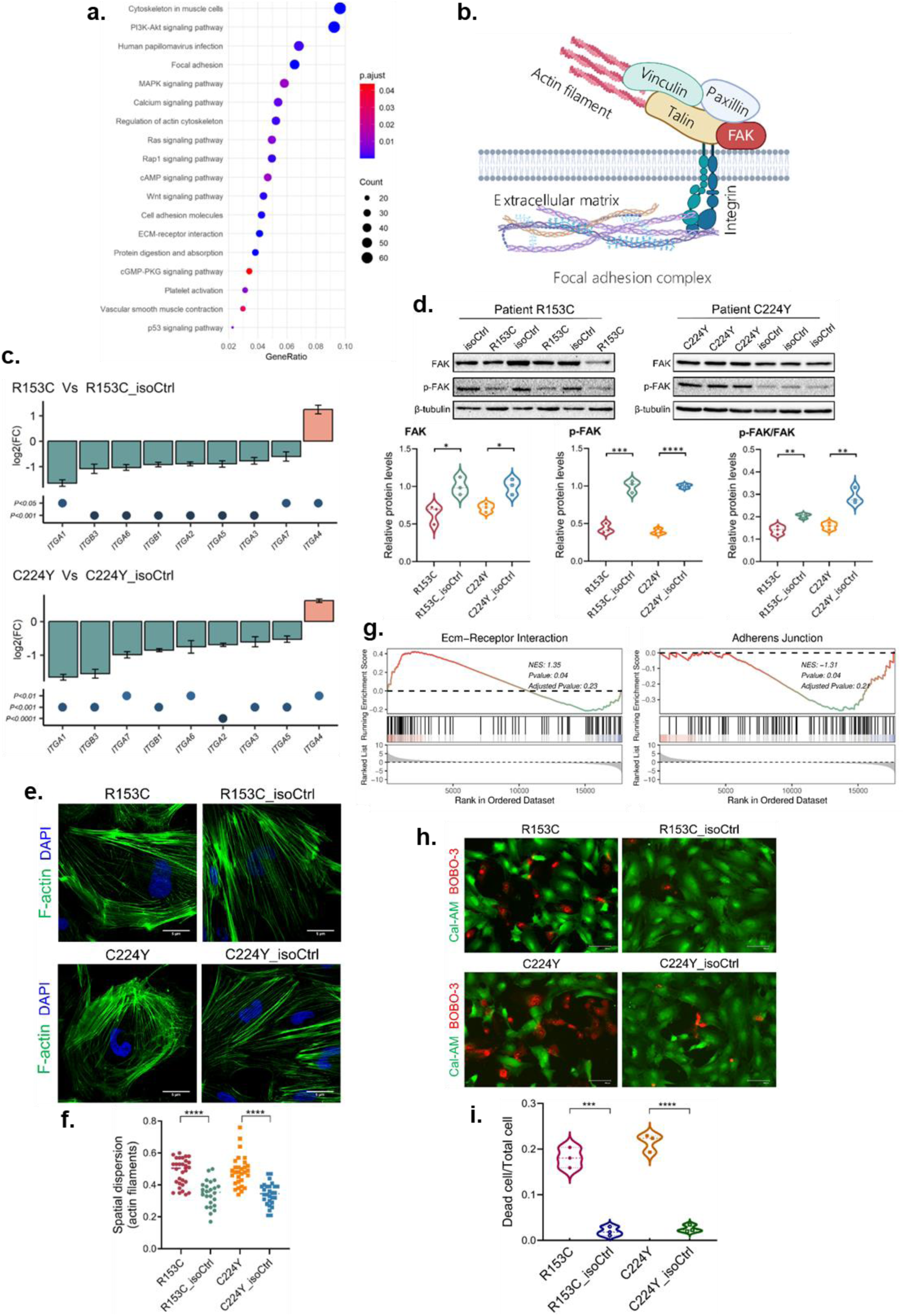
Disruption of cell-matrix and cell-cell interaction of CADASIL iVSMCs leading to cell death. **a** KEGG pathway enrichment of differentially expressed genes (DEGs) between iVSMCs of CADASIL patients and isogenic controls on the RNA sequencing data. **b** A schematic diagram illustrating main components of focal adhesion complex. **c** RT-qPCR determination of the expression of integrin subtypes in iVSMCs derived from iPSCs of CADASIL and their isogenic controls (isoCtrls). Data are presented as mean ± SEM (n=3). Fold changes of gene expression for each gene were calculated separately and compared by unpaired Student’s *t* test. **d** Western blotting of (WB) FAK and p-FAK in iVSMCs derived from iPSCs of CADASIL and isoCtrl. Quantifications of the WB results are shown in the lower panel. Data are presented as mean ± SEM (n=3). Unpaired Student’s *t* test, *p ≤ 0.05, **p ≤ 0.01, ***p ≤ 0.001. **e** F-actin staining of iVSMCs derived from iPSCs of CADASIL and isoCtrls. Nuclei were counterstained by DAPI. Scale bar = 5μm.**f** Organisations of actin cytoskeletons in **e** were analysed by an artificial intelligent (AI) assisted algorithm and presented as special dispersion of actin filament, which was quantified. Unpaired Student’s *t* test, ****p ≤ 0.0001. **g** Gene set enrichment analysis (GSEA) plot of RNA sequencing data on gene Ontology (GO) categories of ECM-receptor interaction and adherens junction in CADASIL iVSMCs versus isoCtrls. **h** Live/dead staining of iVSMCs derived from iPSCs of CADASIL and isoCtrls using Calcein-AM (Cal-AM) and BOBO-3 iodide (BOBO-3), respectively. Scale bar = 100 μm. **i** Quantification of the live/dead staining result in **h**. Data are mean ± SEM (n=3). Unpaired Student’s *t* test, ***p ≤ 0.001, ****p ≤ 0.0001. Results in **c**, **d**, **e** and **h** were all from at least 3 independent iPSC differentiations. Diagram **b** was Created with BioRender.com.

### Nitric oxide signalling is impaired in CADASIL iVSMCs

KEGG analysis of the RNAseq data highlighted the “cGMP-PKG signalling pathway” (**Fig. 6a**). Given the fact that we had already observed reduced *GUCY1A1* (guanylate cyclase 1 subunit alpha 1 / GCSα1) and *GUCY1B1* (guanylate cyclase 1 subunit beta 1 / sGCβ1) in CADASIL iVSMCs (**Fig. 7b-c**) before the RNAseq screening and the prior knowledge of the profound effects of NO in the regulation of VSMC contractility, proliferation, migration, ECM synthesis, and phenotype switching^58–60^, we further explored the involvement of the NO-sGC-cGMP-PKG signalling (**Fig. 7a**) in VSMC pathology of CADASIL. We detected considerable levels of NO in the control iVSMCs with the absence of endothelial cells as measured by DAF-AM staining, which were significantly reduced in the CADASIL iVSMCs (**Fig. 7d**). GCSα1 and GCSβ1 are the two subunits of soluble guanylate cyclase (sGC) that is the primary receptor of NO, which catalyse the synthesis of second messenger cGMP that activate protein kinase G (PKG) inducing VSMC dilation largely via vasodilator-stimulated phosphoprotein (VASP) (**Fig. 7a**)^61,62^. The expression of *PRKG1* and *VASP* and the protein level of VASP were significantly downregulated in CADASIL iVSMCs (**Fig. 7e-f**). To understand the mechanism underlying the reduced NO in CADASIL iVSMCs, we found that endothelial nitric oxide synthase (eNOS) and AKT, a crucial activator of eNOS, were significantly reduced in CADASIL iVSMCs (**Fig. 7g**), while the inducible NOS (iNOS) remained unchanged (**Fig. S8**). eNOS is primarily localised in the plasma membrane and Golgi apparitors, which plays an important role in regulating its enzymatic activity and the bioavailability of NO^63^. We performed immunofluorescence staining and found strikingly that location patterns of eNOS were very different between the CADASIL and isoCtrl iVSMCs. The eNOS signals in the control cells seem associated with intracellular membrane structures, patterns of Golgi and endoplasmic reticulum (ER), whereas the signals in the mutant cells were very weak and defused in cytosol (**Fig. 7h**). As Golgi localisation of eNOS is essential for its posttranslational modifications and activation, the finding likely underlies the compromised NO production (**Fig. 7d**) in CADASIL iVSMCs. Taken together, our results suggest the importance of a local NO-sGC-cGMP signalling in VSMCs independent of ECs, which was dysregulated in CADASIL iVSMCs.

**Figure 7.**
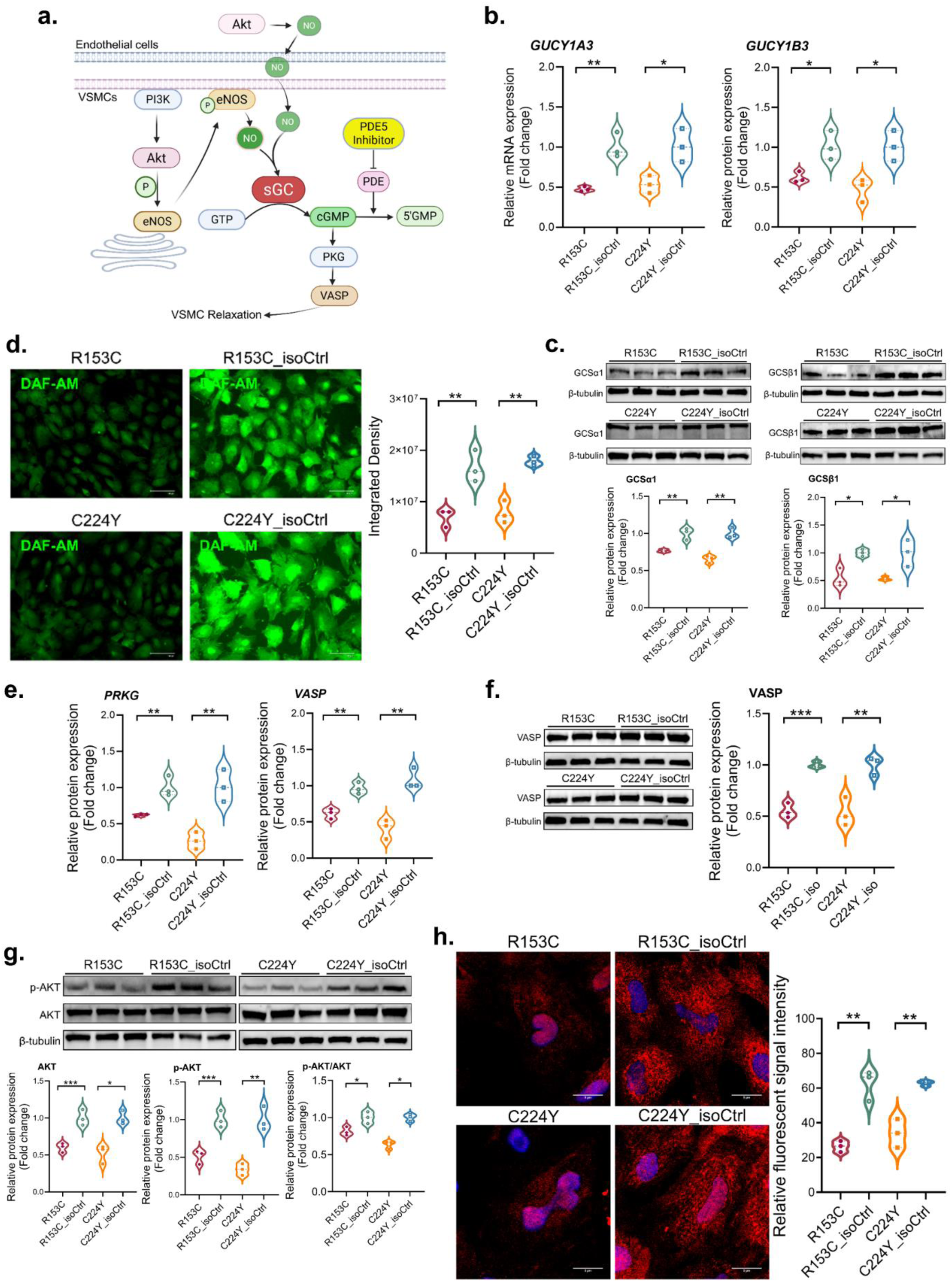
Nitric oxide signalling from iPSC derived VSMCs. IPSCs from two CADASIL patients (R135C & C224Y) and their isogenic controls (R153C_isoCtrl and C224Y_isoCtrl) were differentiated into iVSMCs via neural crest (NC) lineage. **a** Diagram illustrating NO-sGC-cGMP-PKG signalling pathway. **b** & **c** Determination of *GUCY1A1* and *GUCY1B1* expressions by RT-qPCR (**b**) and their protein levels (GCSα1 and GCSβ1) by western blotting (**c**) in iVSMCs derived from CADASIL and isogenic control (isoCtrl) iPSC lines. **d** Cellular nitric oxide (NO) levels in iVSMCs determined by DAF-AM staining and their quantifications (right panel). Scale bar = 100μm. **e** Expressions of *PRKG* and *VASP* in iVSMCs determined by RT-qPCR in iVSMCs. **f** & **g** Western blotting determination of protein levels of VASP (**f**) and AKT and p-AKT (**g**) in iVSMCs. **h** Immunofluorescent staining of eNOS in iVSMCs. Scale bar = 5μm. All quantitative data are presented as mean ± SEM from 3 independent iPSC differentiations (n=3). Unpaired Student’s *t* test, *p ≤ 0.05, **p ≤ 0.01, and ***p ≤ 0.001. Diagram **a** was Created with BioRender.com.

### Treatment of iPSC CADASIL models by PDE5 inhibitor and NO donor significantly restored VSMC function

Building upon our findings, we explored therapeutic potentials by boosting the NO-sGC-cGMP signalling pathway to rescue the key pathological phenotypes of CADASIL iVSMCs. As proof-of-concept experiments, we first supplemented S-nitroso-N-acetylpenicillamine (SNAP), a synthetic NO donor that releases NO under physiological conditions, in cell culture and found that it significantly reversed the abnormal proliferation of CADASIL iVSMCs (**Fig. 8a, 8b** & **S9a**) and moralised protein levels of GCSα1 and GCSβ1 (**Fig. 8d**). However, NO donors usually have short half-life and more suitable in acute settings. To achieve prolonged activation of NO signalling to benefit CADASIL patients and also based on the significant but mild reduction of cGC (*GUCY1A1* and *GUCY1B1*, **Fig. 7c**) in CADASIL iVSMCs, we applied the FDA approved PDE5 inhibitor, sildenafil, that prevents the breakdown of sGC-generated cGMP and activates endogenous sGC-cGMP signalling^64^. Results showed that sildenafil successfully reversed the hyperproliferation of CADASIL iVSMCs to the level of the isogenic controls (**Fig. 8a, 8c** & **S9a**). The treatments also significantly increased the expression of contractile markers α-SMA, calponin and MLCK in the CADASIL iVSMCs (**Fig. 8e**). Interestingly, the disorganised actin cytoskeleton in CADASIL iVSMCs were significantly normalised by both SNAP and sildenafil treatments (**Fig. 8f** & **S9b**). Most importantly, both SNAP and sildenafil successfully prevented the cell death of the CADASIL iVSMCs (**Fig. 8g** & **S9c**).

**Figure 8.**
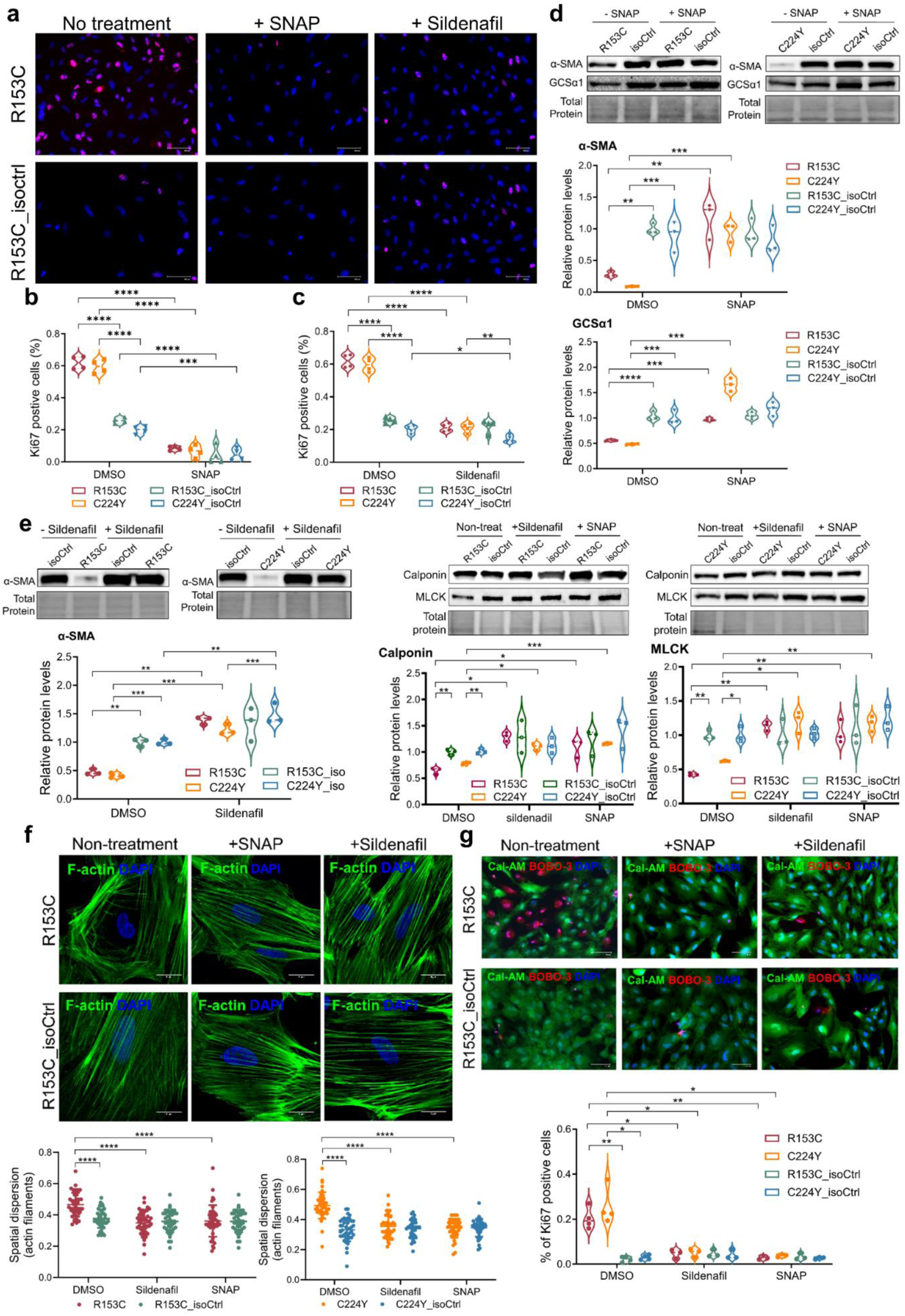
SNAP and Sildenafil rescue of abnormal proliferation, contraction and cell death of CADASIL iVSMCs. IPSCs from two CADASIL patients (R135C & C224Y) and their isogenic controls (R153C_isoCtrl and C224Y_isoCtrl) were differentiated into iVSMCs via neural crest (NC) lineage. The cells were subjected to proliferation assay by Ki67 immunofluorescent staining (**a**) in the presence or absence of the nitric oxide (NO) donor (SNAP) or PDE5 inhibitor (Sildenafil), and results were quantified (**b** & **c**). Scale bars = 100 μm. **d** & **e** Western blotting determination of protein levels of αSMA and GCSα1 with or without SNAP treatment (**d**), αSMA with or without Sildenafil treatment (**e**), and calponin and MLCK with or without SNAP or Sildenafil treatment, respectively, in iVSMCs. Results were quantified and normalised to the total protein loading control (representative areas are shown under each blot). **f** F-actin staining of iVSMCs derived from iPSCs of CADASIL and isoCtrls treated with or without SNAP or Sildenafil. Nuclei were counterstained by DAPI. The organisations of actin cytoskeletons were analysed and quantified by an artificial intelligent (AI) assisted algorithm and presented as special dispersion of actin filament showing in the lower panel. Scale bars = 5 μm. **g** Live/dead staining of iVSMCs derived from iPSCs of CADASIL and isoCtrls using Calcein-AM (Cal-AM) and BOBO-3 iodide (BOBO-3), respectively. Data were quantified shown in the lower panel. Scale bars = 100 μm. All quantitative data are presented as mean ± SEM from 3 independent iPSC differentiations (n=3). Two-way ANOVA and Tukey’s post hoc test, *p ≤ 0.05, **p ≤ 0.01, ***p ≤ 0.001, and ****p ≤ 0.0001.

We previously reported that the CADASIL iVSMCs failed to stabilise the capillary tubule structure formed by ECs with a mechanisms of reduced VEGF secretion form the mutant VSMCs^42^. Considering the role of NO in promoting VEGF synthesis in VSMCs ^65^, we applied SNAP and sildenafil, respectively, during *in vitro* angiogenesis assay, which significantly prolonged the EC tubules stability (**Fig. 9, S10** & **S11**). Our results suggest a promising treatment of CADASIL patients by PDE5 inhibitors to rescue VSMC function and prevent their degeneration seen in CADASIL small vessels.

**Figure 9.**
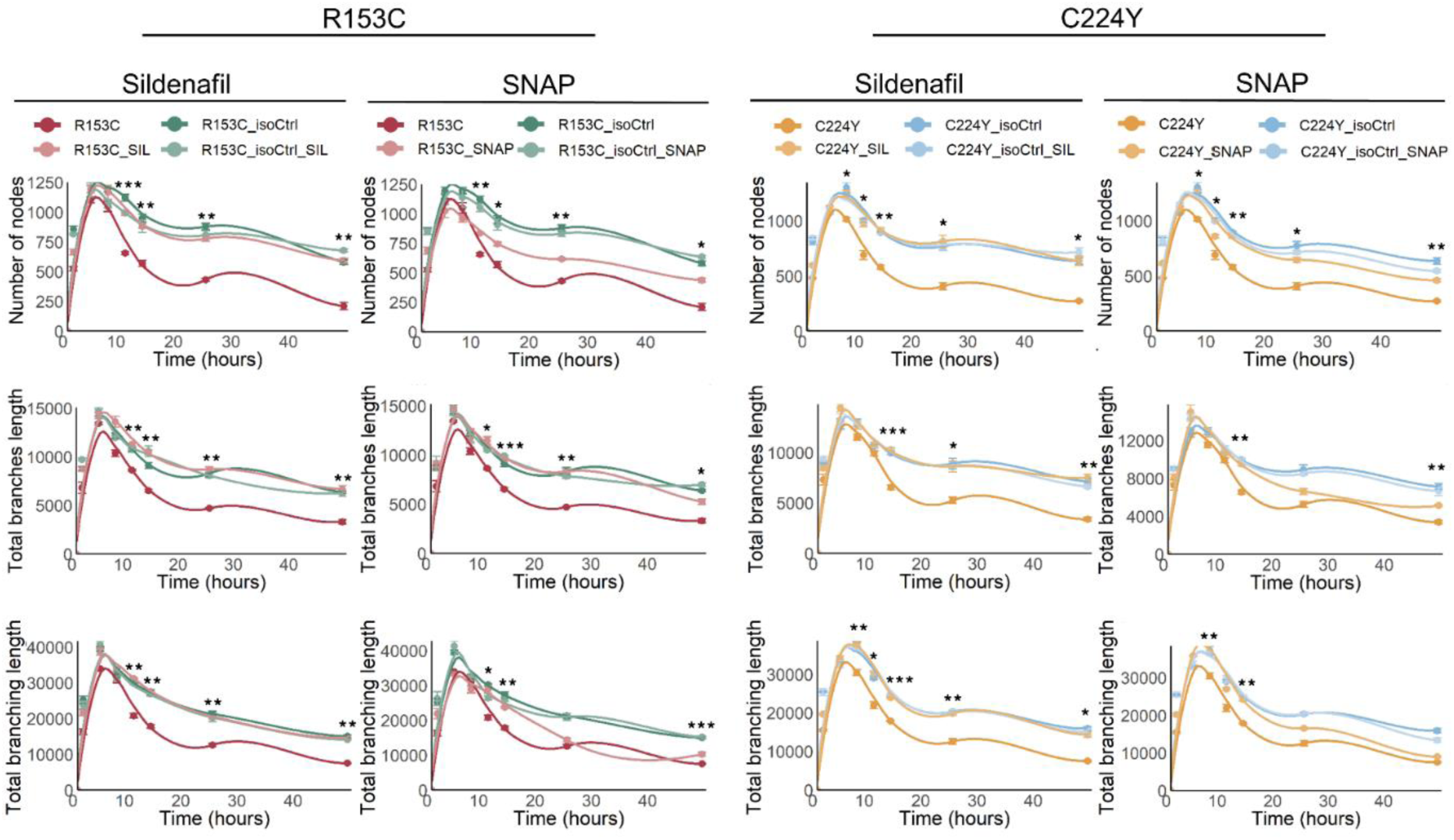
SNAP and Sildenafil rescue of impaired CADASIL iVSMCs in supporting angiogenesis. IPSCs from two CADASIL patients (R135C & C224Y) and their isogenic controls (R153C_isoCtrl and C224Y_isoCtrl) were differentiated into iVSMCs via neural crest lineage. *In vitro* angiogenesis with combined HUVECs and iVSMCs was carried out on Matrigel in the presence or absence of nitric oxide (NO) donor (SNAP) or PDE5 inhibitor (Sildenafil). Quantification of the angiogenesis results showing on the fitted smooth trend lines (LOESS/liner) with 95% confidence interval showing that both SNAP and Sildenafil significantly reversed the impaired ability of CADASIL iVSMCs in supporting the stability of the angiogenic tubule structures. Data are presented as mean ± SEM from 3 independent iPSC differentiations (n=3). Two-way ANOVA and Tukey’s post hoc test were conducted. Asterisk stars (*s) showing significant differences of the treated versus non-treated samples of CADASIL iVSMCs at the indicated time points. *p ≤ 0.05, **p ≤ 0.01, and ***p ≤ 0.001.

## Discussion

Using an iPSC model of CADASIL, we uncovered for the first time that iPSC derived VSMCs via a developmental pathway mimicking VSMCs of cerebral vasculature exhibited a unique susceptibility to CADASIL-associated *NOTCH3* mutations, comparing to iPSC-SMCs that mimic sources of peripheral vasculatures. The CADASIL iVSMCs displayed hyperproliferation, hypermigration, reduced contractility, and abnormal cell-matrix interaction. This study also provided the first comprehensive and systemic evaluation of VSMC phenotype in CADASIL, revealing a switching of CADASIL iVSMCs from a contractile to a synthetic phenotype and reinforcing the role of ECM disorganisation in CADASIL pathology. Additionally, we provided evidence of disrupted cell-matrix and cell-cell adhesion of CADASIL iVSMCs, which likely leads to a loss of cell anchorage and subsequent *anoikis*. Finally, alterations in the VSMC-specific NO-sGC-cGMP signalling and phenotype rescue of CADASIL iVSMCs by PDE5 inhibitor, sildenafil, highlight a potential novel therapeutic avenue for CADASIL.

It has been intriguing that CADASIL, a systemic vasculopathy, has prominent manifestations in the brain leading to strokes and vascular dementia while rarely primarily presented as myocardial infarction or symptoms of peripheral vessel disease. Our findings revealed a selective vulnerability of the CADASIL NC-SMCs compared to the LPM-SMCs and PM-SMCs, offering a potential explanation of the paradox. Developmental origin-specific VSMCs exhibit distinct functionalities, including differences in growth behaviours and responses to stimuli^28–30,48,49,66,67^. These differences are intrinsic and driven by unique gene expression profiles including the differential expressed *HOX* genes in lineage-specific VSMCs^47,68–70^. For example, in a mouse model, when vessels prone to atherosclerosis were grafted to low-pressure area of the vascular tree, they developed atherosclerosis in the absence of mechanical and haemodynamic factors^71^, highlighting a location-specific tissue susceptibility to disease. Different reactivities of VSMC subtypes to stimulations by growth factors or cytokines likely underlie different disease susceptibility in vessel beds of different locations. Angiotensin II was found to promote proliferation of VSMCs isolated from ascending aorta but not from other aorta regions, whereas PDGF-BB promoted proliferation in VSMCs from all aortic regions^29^. Additionally, TGF-β1 preferentially stimulated the growth and proliferation of neural crest derived VSMCs over mesoderm derived VSMCs via c-myb^30,66^, and interestingly, the former resulted in marked elevation of α1 collagen and the latter had no effect on the α1 collagen gene expression^30^. Based on these observations, it is not surprising to see a selective susceptibility of the cerebral VSMCs to the *NOTCH3* mutations in CADASIL. Our finding is in line with a pathological report on a postmortem CADASIL patient where minimal changes were observed in liver, spleen, heart and lung whereas significant VSMC disruptions were observed in all small arterioles and capillaries in the brain sample^54^.

It is worth noting that what we identified was the vulnerability of the brain VSMCs to CADASIL *NOTCH3* variants which altered key functionalities of the mutant VSMCs, i.e., proliferation, migration, cell-matrix interaction, and contraction. This does not exclude the possibility of peripheral VSMCs developing pathological changes via alternative mechanisms or in a later stage. This also should not override the systemic GOM deposition and Notch3 accumulation that is largely attribute to the conformational change of mutant Notch3 proteins due to cysteine alterations^72^. Nevertheless, the increased sensitivity of brain VSMCs to *NOTCH3* variants likely accelerate the CNS phenotype in CADASIL, suggesting early intervention could benefit CADASIL patients. As our experiments were mostly conducted on 2D monolayer cell cultures, which is difficult for GOM or Notch3 to accumulate, it is therefore attempt to speculate that functional changes reported in isolated primary peripheral VSMCs from CADASIL patients or animal models could mainly attribute to protein accumulations that are systemic. It would be interesting to clarify this as it has not been convincing evidence on whether GOM or Notch3 accumulation are the primary driver of the CADASIL pathology or a secondary effect or readout of the pathology. Overall, our finding contributes to refine future therapeutic targets to cerebral vasculatures for CADASIL and also highlights the importance of using correct models in studying human disease with iPSCs. Future study is to clarify the molecular mechanisms by which *NOTCH3* variants specifically alter the functionality of cerebral VSMCs.

Findings from previous studies on increased proliferation of VSMCs from CADASIL patients^39^ and matrix deposition in small arteries of CADASIL mice^73^ suggested a phenotype change of VSMCs. However, there have not been studies specifically addressing the VSMC phenotype in CADASIL. Through a comprehensive analysis of transcriptomic data, RT-qPCR, western blotting, immunofluorescent images, and functional assays, we concluded that CADASIL iPSC-SMCs switched from a contractile to synthetic phenotype. The CADASIL iVSMCs highly expressed a set of synthetic markers, decreased expression of contractile related genes, as well as reduced contractility and enhanced proliferation and migration, compared to isogenic controls. Synthetic VSMCs exhibit a secretory feature, which would contribute to the ECM accumulation found in CADASIL arteries. However, the accumulated and disorganised ECM may play an even significant role in driving VSMCs to switch their phenotype. ECM is equally important as growth factors in regulating VSMC phenotypes through binding to their integrin receptors^36,74^. We have found downregulation of a range of integrin genes expressed in the CADASIL iVSMCs, which likely contributed to the VSMC phenotype change. We believe that a vicious cycle is formed where synthetic VSMCs abnormally produce matrix whereas accumulated matrix drives VSMC phenotype changes. Since phenotype switching is a hallmark of atherosclerosis, a target for actively developing therapeutic approaches including those involving non-coding RNAs^75,76^, highlighting VSMC phenotype switching in CADASIL could provide opportunities to repurpose treatments originally developed for more common conditions.

It was observed in early year CADASIL research that VSMCs from both human CADASIL patients or mouse models lost anchorage to adjacent cells and ECM, from which it was hypothesised that this might be one of the key events initiating the cascade leading to VSMC degeneration in CADASIL^54,77^. In light of the essential role of integrin-ECM engagement in generating pro-survival signals through activation of FAK and the subsequent recruitment of vinculin and the focal adhesion formation^78–82^, our finding of the significant reduction of FAK, pFAK, and integrins as well as disorganised vinculin in CADASIL iVSMCs provided molecular basis supporting the hypothesis. GESA enriched “adherence junctions” and “cell junction organisation” as well as TEM imaging further support the loss of cell-cell interaction. Abnormal cell-matrix and cell-cell adhesions usually lead to *anoikis*, a type of apoptosis induced by cell detachment^56,78,83^. Indeed, we observed significant level of cell death (**Fig. 6h**) and increased apoptosis in our previous finding^42^. Thus, the CADASIL iPSCs could be a useful human model for future drug discovery aiming to reverse VSMC survival in this condition.

We provided several lines of evidence demonstrating a significant impairment of NO-sGC-cGMP-PKG pathway in CADASIL iVSMCs (**Fig. 7**). For the first time we observed a reduced NO level in the CADASIL iVSMCs. This highlighted a NO-sGC-cGMP-PKG signalling pathway in VSMCs ^84–86^ with the absence of ECs that usually provide NO for VSMCs for vessel dilatation. While Akt is usually responsible for the phosphorylation and activation of eNOS in ECs, but we identified for the first time that Akt and pAkt were significantly reduced in the CADASIL iVSMCs, which likely contributes to the reduction of NO. Translocation to the Golgi apparatus is required for eNOS palmitoylation, which is essential for eNOS to recycle back to cell membrane caveolae to be activated by Akt^63^. The abnormal location and reduced level of eNOS could affect its activation further reducing NO levels in the CADASIL iVSMCs. Consequently, the NO receptor, sGC subunits, were downregulated at both gene (*GUCY1A1* and *GUCY1B1*) and protein levels, and cGMP effector PKG and VASP were reduced. However, the reduced expression of *GUCY1A1* and *GUCY1B1* could via other mechanisms. It has previously reported an RBP-Jκ binding site on the *GUCY1A3* promoter region in the developing heart^87^, linking the canonical Notch signalling to the regulation of sGC. Therefore, the reduced activity of mutant Notch3 (**Fig. S8**), although mild, may partially contribute to the downregulation of *GUCY1A3* in CADASIL VSMCs. Our finding is in line with a previous report where cGMP was reduced in VSMCs from CADASIL mice and patients, although it was caused by oxidation of sGCβ1^88^.

To target the damaged NO-sGC-cGMP signalling pathway, we demonstrated that both NO donor SNAP and PDE5 inhibitor Sildenafil could effectively rescue the expression of contractile markers, the abnormal proliferation and angiogenesis, and cell death in the CADASIL iVSMCs. PDE5 specifically targets cellular cGMP for its degradation^64^. Sildenafil, an FDA approved medicine primarily for the treatment of erectile dysfunction, is a PDE5-specific inhibitor augmenting the NO-sGC-cGMP signalling. Although there are controversial findings about the effect of NO on VSMC proliferation, migration and phenotype switching^59^, general understanding support a protective role of NO on VSMCs behaviours and functions^58,60,89^. Sildenafil also rescued the impaired function of iVSMCs in stabilising angiogenesis structures which was found in our previous report involving reduced VEGF secretion from the mutant mural cells^42^. It is already known that NO induces the synthesis of VEGF in VSMCs^65^ and endogenous NO from endothelial cells promoted angiogenesis^90^. Interestingly, sildenafil has already been found to promote angiogenesis ^64,91,92^ and upregulate VEGF^93^. All these findings support our results.

To date, accumulating evidence has demonstrated beneficial effects of PDE5 inhibitors in the treatment of a range common conditions including myocardial infarction, cardiac hypertrophy, heart failure, diabetes, cancer, Duchenne muscular dystrophy, Alzheimer’s disease, and aging-related conditions^64^. Most interestingly, a recent work based on a retrospective case-control pharmacoepidemiologic analyses of insurance claims data that included 7.23 million individuals revealed that sildenafil usage was associated with a 69% reduction of Alzheimer’s disease risk^94^. This finding is highly encouraging in relation to our findings in the vascular dementia CADASIL. It has been found that PDE5 exists not only in vascular cells but also in human neurons, which provide a viable target for neurologic disease^90^. Indeed, PDE5 inhibition rescued the oligomeric form of tau protein induced reduction of long-term potentiation and memory impairment^64,95^. New PDE5 inhibitors have been actively developing with a hope to improve its potency, selectivity and BBB permeability^64,96^. The beneficial roles of PDE5 inhibition on both vascular and neurologic functions make it a good candidate for vascular dementia. Therefore, our finding represents a novel but viable treatment for CADASIL, which likely benefit from the drug development for Alzheimer’s disease in this area in the future.

Recently, the U.S. Food and Drug Administration (FDA) passed the Modernization Act 3.0, which eliminated the mandate requirement for animal testing and allowed human cell models to be used in preclinical drug development^97,98^. This not only reduces the reliance on animal experimentation, but also greatly encourages the use of iPSC models in human disease research. Our study demonstrated a robust iPSC CADASIL model, which can be used to further investigate disease mechanisms and facilitate drug development, ultimately benefit patients.

## Methods

### Cell culture

The human induced pluripotent stem cell (iPSC) lines derived from two CADASIL patients carrying the *NOTCH3* heterozygous variants R153C and C224Y and healthy controls, respectively, were reported in our previous studies with local ethical approval (REC reference no. 12/NW/0533)^42,43^. Isogenic control lines (R153C_iso and C224Y_iso) were created using CRISPR/Cas9 gene editing. IPSCs were maintained in Essential 8 medium (Gibco; A1517001) in CO_2_ incubator at 37°C. For passaging, sub-confluent iPSCs were dissociated using 0.5 mM EDTA (Invitrogen; AM9260G) in PBS at 37 °C for 5–6 minutes. Cells were gently rinsed by pipetting in 1 mL of Essential 8 medium, yielding small aggregates which were seeded onto vitronectin (VTN; 0.005mg/ml; Gibco; A14700)-coated plates at a density of 2.5–3 × 10³ cells/cm² in Essential 8 medium supplemented with 10 µM Y-27632 (Tocris; 1254). After 24 hours, the medium was replaced with fresh Essential 8 medium without Y-27632.

### Lineage-specific vascular smooth muscle cell (VSMC) differentiation from iPSCs

Differentiation of VSMCs from iPSCs were performed according to published methods^32,99^ with minor modifications.

#### Neural crest cell differentiation

To generate neural crest progenitor cells, iPSCs were cultured on VTN-coated plates in chemically defined Essential 6 medium (Gibco; A1516401) with 10 ng/mL FGF2 (PeproTech; 100-18B) and 10 μM SB431542 (Tocris; 1614). Cells were first passaged at day 5 and subsequently upon reaching confluence, with continued supplementation of FGF2 and SB431542, up to passage 12.

#### Mesodermal progenitor cell differentiation

To induce early mesoderm, human iPSCs were cultured in Essential 6 medium supplemented with 20 ng/mL FGF2, 10 uM LY294002 (Selleckchem; S1105) and 25 ng/mL BMP4 (PeproTech; 120-05ET) for 48 hours. Subsequent mesodermal subtype specification was directed over the following 4 days. Lateral plate mesoderm (LPM) differentiation was achieved by continued exposure to 20 ng/mL FGF2 and 50 ng/mL BMP4, while paraxial mesoderm (PM) differentiation was promoted using 20 ng/mL FGF2 and 10 µM LY294002.

#### VSMC differentiation from neural crest cells and mesodermal progenitors

Following the generation of intermediate populations, cells were trypsinized and cultured in SMC differentiation medium composed of Essential 6 supplemented with 10 ng/mL PDGF-BB (PeproTech; 100-14B) and 2 ng/mL TGF-β1 (PeproTech; 100-21) for a minimum of 12 days. Neural crest-derived VSMCs (NC-SMCs) reached maturity by day18, while LPM- and PM-derived SMCs matured by day 16. These mature iPSC-derived lineage specific VSMCs (iPSC-VSMCs) were subsequently maintained in Smooth Muscle Cell Growth Medium 2 (SMCG2; PromoCell; C-22062) for long-term culture, up to 5 passages.

### Cell proliferation analysis

Cell proliferation was evaluated by Ki67 immunostaining assay. Mature iPSC-VSMCs were seeded on VTN-coated 24-swell plates at Day 16 for NC-SMCs and Day14 for LPM- and PM-SMCs. After 48 hours interval, cells were fixed with 4% paraformaldehyde (PFA; Thermo Fisher; J61899.AK) and subjected to immunostaining for the proliferation marker Ki67 (Abcam; ab15580). The percentage of Ki67-positive cells was quantified using ImageJ software, based on analysis of at least six randomly selected fields per condition in each experiment. The experiment was independently repeated using iPSC-VSMCs derived from three separate rounds of differentiation.

### Cell migration assay

The migratory behaviour of iPSC-VSMCs was assessed by the wound healing assay using the IncuCyte live-cell imaging system (Sartorius; Essen Bioscience). Mature iPSC-VSMCs (4X10^4^ cells/well) were seeded onto VTN-coated 96-well ImageLock tissue culture plate (Essen Bioscience) and incubated at 37 °C with 5% CO₂ 24 hours to allow cell attachment and confluence. Wounds were introduced by 96-well WoundMaker (Essen Bioscience), followed by two gentle PBS washes to remove detached cells. Wound closure was monitored in real-time by capturing images at 2-hour intervals. Migration was quantified based on changes in wound confluence over time using the IncuCyte analysis software (Essen BioScience). Each condition was tested in technical duplicate, and the experiment was independently repeated using iPSC-VSMCs derived from three separate differentiations

### Collagen-I contraction assay

Contractile function of iPSC-VSMCs was investigated by collagen I gel contraction assay. The mature iPSC-VSMCs (1X10^5^ cells/well) were re-suspended and mixed with 150μL of PureCol collagen solution (Advanced Biomatrix; 5005) per well. The collagen-cell suspension was dispensed into 48-well tissue culture plates and allowed to polymerise for an hour at 37 °C. After polymerisation, the gels were mechanically released by gently running a sterile spatula around the well perimeter, followed by a brief flush with medium to ensure complete detachment. Gel contraction was monitored, and images were captured at 48 hours using a ChemiDoc XRS imaging system (Bio-Rad). The area of each gel was quantified using ImageJ software to assess contraction. For each condition, three technical replicates were included, and the experiment was repeated three times using iPSC-VSMCs derived from independent differentiation batches.

### Measurement of intracellular NO levels

Intracellular nitric oxide (NO) levels in NC-SMCs derived from CADASIL patients and isoCtrls was measured using fluorescent probe 4-amino-5-methylamino-2′,7′-difluorofluorescein diacetate (DAF-FM; Invitrogen; D23844). Mature NC-SMCs were incubated with 10 μM DAF-FM diacetate in cell culture medium for 30 min at 37 °. Following probe loading, cells were washed with PBS and incubated in fresh medium for another 30 min to allow complete intracellular de-esterification. Fluorescence was visualised using an EVOS M3000 microscope with 488 nm excitation. Quantification of fluorescence intensity was performed using ImageJ, with integrated density measurements was quantified using ImageJ. Images were acquired from five randomly selected fields per condition, and experiments were conducted in triplicate using NC-SMCs obtained from three independent differentiations.

### Cell viability assay

Cell survival of NC-SMCs from CADASIL patients, isoCtrls, and those treated with activators of the NO–cGMP–PKG signalling pathway was assessed using the LIVE/DEAD Cell Imaging Kit (488/570; Invitrogen; R37601). Mature NC-SMCs were cultured in 24-well plate and cultured for 48 hours at 37 °C in a 5% incubator to make cell recovery. Cells were stained according to the manufacturer’s instructions, together with counterstaining Hoechst 33342 (300ng/ml) to label the nuclei. Fluorescent images were acquired using the EVOS M3000 imaging system (Thermo Fisher) with excitation at 488 nm (live cells) and 570 nm (dead cells). Cell viability was calculated as the ratio of live cells to total cells (live + dead) based on Hoechst-positive nuclei. For each condition, five random fields were imaged per well, and the experiment was independently repeated three times using NC-SMCs derived from three separate rounds of differentiation.

### Angiogenesis assay

The angiogenic response of NC-SMCs from CADASIL patients, isoCtrls, and those treated with activators of the NO–cGMP–PKG signalling pathway was assessed using Matrigel-based endothelial tube formation assay. Growth factor-reduced Matrigel (Corning; 354230) was thawed on ice, dispensed (50 μl/well) into pre-chilled 96-well plates, and incubated overnight at 37 °C in a 5% incubator to allow polymerization. A co-culture of 1.5×10^4^ human umbilical vein endothelial cells (HUVECs) and 0.75 ×10^4^ NC-SMCs was plated onto the thin layer of Matrigel in essential 6 supplemented with 5ng/ml VEGF-165 (PeproTech; 100-20) and 2ng/ml FGF_2_. Cultures were maintained at 37 °C in a humidified 5% CO_2_ incubator to promote capillary-like network formation. Tube-like structures were captured at 1, 4, 7, 10, 13, 24, and 48 hours using Brightfield microscopy (EVOS; Thermo Fisher). Quantification of angiogenic parameters, including the number of nodes, total branching length, and total branches length, was performed using imageJ with the “Angiogenesis Analyser” plugin. Each condition was test in technical duplicates, and the experiment was repeated independently three times using iVSMCs derived from three separate rounds of differentiation.

### Generation of spheroids

Size-matched spheroids were generated by Day 18 NC-SMCs derived from CADASIL patients and isoCtrls using the hanging drop method. Briefly, cells were dissociated with trypsin-EDTA, centrifuged at 300Xg for 5 minutes, re-suspended and counted using an automated cell counter. Approximately 1X10^5^ cells in 20 uL of SMCG2 medium supplemented with 4% poly(vinyl alcohol) (PVA; Sigma-Aldrich; 341584) were seeded per drop. After 24 hours, spheroids were collected and transferred to low attachment 96-well plates. The SMCG2 medium was first replaced at 24 h and subsequently every 48 h for the duration of the culture.

### 3D migration assay

The migratory capacity of NC-SMCs derived from CADASIL patients and isoCtrls was evaluated by 3D spheroid migration assay. For the 3D migration assay, spheroids were cultured for 24 hours and were subsequently embedded in collagen I. The collagen solution was prepared by mixing collagen I matrices (Advanced Biomatrix; 5005) with cell culture medium at a 2:1 ratio, and approximately 300 µL of the mixture was added per well in a 48-well plate. After a 5-minute settling period, 4–5 spheroids were randomly placed into each well. Plates were incubated at 37 °C with 5% CO₂ for 1 hour to allow gel polymerization. To inhibit proliferation and focus on cell migration, spheroids were treated with 25 µg/mL mitomycin C (Stemcell Technology; 73274) for 20 min, followed by replacement with fresh culture medium. Brightfield images were acquired daily to monitor spheroid outgrowth. Migration distances were quantified using ImageJ by converting images to 8-bit format with the “Mask” function to enhance visualization and measurement. Each data point in the violin plot represents an individual spheroid. Spheroids were aggregated from NC-SMCs derived from three independent differentiations for biological repeats.

### Transmission Electron Microscope (TEM) protocol for spheroids

Spheroids were fixed immediately in a solution of 2.5% glutaraldehyde and 4% formaldehyde in 0.1 M HEPES buffer, pH 7.2 2 h. Samples were then rinsed in distilled water. Samples were then post-fixed in a solution of 1.0% osmium tetroxide and 1.5% potassium ferrocyanide in 0.1M cacodylate buffer for 2 h and washed with H2O for 3 times to remove osmium tetroxide residuals. Specimens underwent incubation in thiocarbohydrazide (TCH) for 60 min at RT followed by 5 times washing with distilled water, until any formed TCH crystals are dissolved. Then these spheroids were incubated in 1% osmium tetroxide in distilled water for 1 hour before incubated in 1% uranyl acetate for another an hour.The these specimen underwent dehydration steps in ascending ethanol series (30%, 50 %, 70%, 90%, 100% v/v × 3). Speciment were infiltrated in graded TAAB 812 Hard in acetone at RT overnight. Finally, samples were embedded in free 100% TAAB 812 Hard in labelled mould and put in a stove at 60 °C for 48 h. Ultrathin sections for transmission electron microscopy observations were cut using an ultramicrotome (Leica EM UC6, Vienna, Austria). Ultrathin sections were collected on 100-mesh copper grids (Assing, Rome, Italy) stained with Uranyless© solution and lead citrate 3% solution (Electron Microscopy Science, 1560 Industry Road, Hatfield, PA, USA). Imaging was performed by a transmission electron microscope (Carl Zeiss EM10, Thornwood, NY, USA) set with an accelerating voltage of 60 kV. Images were acquired with a CCD digital camera (AMT CCD, Deben UK Ltd., Suffolk, UK).

### Luciferase reporter assay

The activation of NOTCH signalling pathway was detected by Dual-Glo Luciferase Assay System (Promega; E2920) as we previously described^21^. The mature NC-SMCs from CADASIL patients and isoCtrls were plated into a 12 well plate and transfected with 4xCSL-luciferase reporter vector (Addgene plasmid 41726,1μg) and Renilla luciferase (0.5μg) using lipofectamine LTX reagent with Plus reagent (Thermo Fisher, 15338030) following the manufacturer’s instructions. After 24 hours of transfection, the activity of firefly luciferase and renilla luciferase were quantified in 96-well plate by a luminometer using Dual-Glo luciferase kit according to the instructions provided by the manufacturer.

### Real-time quantitative polymerase chain reaction

Cells were harvested from 6-well culture plates, and total RNA was extracted using RNeasy Mini kit (Qiagen; 74104). Then Complementary DNA (cDNA) was synthesized using the high-capacity RNA-to-cDNA kit (Invitrogen; 4388950). Quantitative real-time PCR was performed using the QuantStudio 6 Real-Time system (Thermo Fisher) using Power SYBR Green PCR Master Mix (Applied Biosystems; 4367659). Gene expression levels were normalized to *GAPDH* as the housekeeping gene, and relative quantification was calculated using the Pfaffl method. Each sample was test in technical triplicate, and the experiment was repeated independently three times using iPSC-VSMCs derived from three separate rounds of differentiation. The primer sequences and efficiency are shown in Table S1.

### Western Blotting

Cells were lysed with RIPA buffer (Sigma-Aldrich; R0278) supplemented with 100X EDTA-free Halt Protease Inhibitor Cocktail (Thermo Fisher, 87785), and lysates were centrifuged at 16200 x g for 20 min at 4°C. The supernatant was collected, and protein concentration was determined using the Pierce BCA protein assay kit (Thermo Fisher, 23225). For each sample, 20μg of total protein was mixed with NuPAGE LDS sample buffer and β-mercaptoethanol, then boiled at 95°C for 5 min. Proteins were resolved on 12% or 4-20% TGX stain-free precast gels (Bio-rad) and transferred onto nitrocellulose membranes. Membranes were blocked in TBS containing 0.1% (v/v) Tween-20 (TBST) supplemented with 5% (w/v) slim milk powder and 2% (w/v) BSA for an hour at RT, followed by incubation with primary antibodies overnight at 4°C. After washing with TBST, blots were incubated with proper HRP-conjugated secondary antibodies for 2 hours at RT. Signal detection was performed using SuperSignal™ West Pico or Femto Chemiluminescent Substrate (Thermo Fisher Scientific), and blots were imaged using autoradiography. Where indicated, band intensities were quantified by densitometry using Image Lab software (Bio-Rad).

### Immunofluorescence on 3D spheroids

Immunofluorescence (IF) staining was performed to assess NOTCH3 (R&D; AF1308) and collagen IV (Thermo Fisher; PA5-104508) expression in 3D spheroids derived from NC-SMCs. Spheroids were cultured in SMCG2 medium for an additional 7 days to promote extracellular matrix formation. Subsequently, spheroids were collected into 1.5 ml Eppendorf tubes, washed with PBS, and fixed in 4% PFA for 2 hours at 4°C. Fixed spheroids were permeabilized with 0.5% (v/v) Triton X-100 in PBS for 4 hours at room temperature (RT), followed by blocking overnight in IF buffer containing 10 % donkey serum, 2% (w/v) BSA, 0.2% (v/v) Triton X-100, and 0.05% (v/v) Tween-20 in PBS. Primary anti-human antibodies NOTCH3 and Collagen IV were diluted in fresh IF buffer and incubated the spheroids for 3 days at RT. After primary incubation, spheroids were washed 3 times with IF buffer and incubated for 24 hours at RT with species-specific secondary antibodies (anti-goat Alexa Fluor 488 and anti-rabbit Alexa Fluor 647) and Hoechst 33342 (1:50,000 dilution) diluted in IF buffer. Following three additional washes, spheroids were mounted in 0.5 mm deep iSpacer chambers filled with Rapiclear 1.49 (Sunjin Lab Co.). Spheroids were imaged using a Leica SP8 confocal microscope under a 10x objective. Z-stacks were acquired and processed using ImageJ to generate either 2D maximum intensity projections or 3D reconstructions.

### Immunofluorescence on monolayer iPSC-VSMCs

IF was performed to assess the expression of VSMC-specific or functional markers in iPSC-VSMCs. Cells were cultured on VTN-coated coverslips or 24-well plates and fixed with 4% PFA for 15 minutes at RT. Following fixation, cells were permeabilized with 0.5% Triton X-100 in PBS for 20 min and blocked in IF buffer an hour at RT. Cells were then incubated with primary antibodies: anti-αSMA antibody (1:250, Abcam, ab7817), anti-calponin (1:250, Abcam, ab46794), anti-transgelin (1:500, Abcam, ab14106), anti-vinculin (1:500, Sigma-Aldrich, V9131), anti-eNOS (1:200; Novus, NB300) diluted in IF buffer overnight at RT. After washing, appropriate Alexa Fluor–conjugated secondary antibodies were applied for two hours at room temperature in darkness. Nuclei were counterstained with DAPI. Coverslips were mounted using Fluoromount-G mounting medium (Invitrogen; 00-4958-02), and images were acquired using a fluorescence or Leica SP8 confocal microscope. Quantification of fluorescence intensity or positive cell percentage was performed using ImageJ.

### F-actin staining and quantification

Immature NC-SMCs (Day12) were placed onto VTN-coated glass coverslips and cultured in SMC differentiation medium. At day 18, cells were fixed with 4% PFA for 15 minutes at RT, permeabilized with 0.5% Triton X-100 in PBS for 10 minutes, and blocked in IF buffer for 30 minutes. For visualization of the cytoskeletal structure, F-actin was stained using Phalloidin-iFluor 488 Reagent (Abcam, ab176753) for 2 hours at RT. Nuclei were counterstained with DAPI. Images were acquired using a Leica SP8 inverted confocal microscope equipped with a 100× oil immersion objective.

Spatial dispersion is used to quantify cytoskeletal organization. We resize the input image to 512×512 pixels. Then, the standard Canny edge detection algorithm^100^ is employed to outline the edges of stress fibres. Firstly, a Gaussian filter is applied to smoothing the image whilst reducing image noise. Afterwards, image gradients are computed so that potential edges can be better identified. The detected edges are then processed using a Non-Maximum Suppression (NMS) technique^101^ to eliminate false detections, followed by edge screening via a dual-threshold approach of Canny edge detection^100^. The dual thresholds are determined, based on statistical characteristics of image pixels. To be specific, the standard Sobel operator^102^ is used to calculate horizontal (*x*) and vertical (*y*) gradients (*G*_*x*_ and *G*_*y*_). The gradient magnitude *G* at each pixel, representing gradient intensity, is computed as:

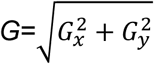

The mean (*M*) and standard deviation (*S*) of the gradient magnitudes are then calculated. The upper threshold (*T*_*max*_) and the lower threshold (*T*_*min*_) are determined as follows, where T denotes the reduction ratio:

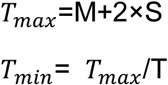

Next, the resized image is converted to the counterpart in the HSV colour space. The centre point of the blue region is identified based on a predefined HSV range. For each stress fibre edge, two parameters are computed:

1. The distance (*D*^*i*^) from the centre of the blue region to the edge.
2. The angle (*A*^*i*^) between the edge and the horizontal axis (0° < *A*^*i*^ < 180°).

For the images containing multiple blue regions, the average distance (*D^*i*^_avg_*) from each stress fibre edge to all the blue centres is calculated individually. During this process, small blue regions and short stress fibre edges are filtered out.

Finally, spatial dispersion (*dsp*) is used to measure the degree of disorder. Each pair (*D^*i*^_avg_*, *A*^*i*^) is treated as a point in the 2D space. The space is divided into *n* subspaces, and the probability *p* of the points falling into each subspace is computed as the ratio of the points in that specific subspace against the total number of the points. The entropy (*etp*) and the normalized spatial dispersion (*dsp*) are derived as follows:

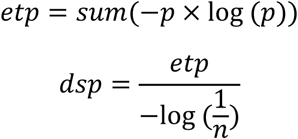

where *etp* is used to measure the uncertainty of the point distribution within spatial sub-regions, where *p* represents the proportion of the points in each sub-region relative to the total number of the points. *n* denotes the total number of the sub-regions. By dividing *etp* by the maximum entropy (log_2_ *n*), the result is normalized to a scale of [0,1], facilitating the comparisons across different regions.

This code is available at https://github.com/zhaoaite/ActinDetection.

### RNA-seq sample processing, sequencing and data processing

Total RNA isolated using the abovementioned method was submitted to the Genomic Technologies Core Facility (GTCF) at the University of Manchester. The quality and integrity of the RNA samples were assessed using a 4200 TapeStation (Agilent Technologies, Cheadle, UK) and then libraries generated using the Illumina® Stranded mRNA Prep. Ligation kit (Illumina, Inc., Cambridge, UK) according to the manufacturer’s protocol. Briefly, total RNA (typically 0.025–1μg) was used as input material, from which polyadenylated mRNA was purified using poly-T, oligo-attached, magnetic beads. Next, the mRNA was fragmented under elevated temperature and then reverse transcribed into first-strand cDNA using random hexamer primers and in the presence of Actinomycin D (thus improving strand specificity whilst mitigating spurious DNA-dependent synthesis). Following removal of the template RNA, second-strand cDNA was then synthesised to yield blunt-ended, double-stranded cDNA fragments. Strand specificity was maintained by the incorporation of deoxyuridine triphosphate (dUTP) in place of dTTP to quench the second strand during subsequent amplification. Following a single adenine (A) base addition, adapters with a corresponding, complementary thymine (T) overhang were ligated to the cDNA fragments. Pre-index anchors were then ligated to the ends of the double-stranded cDNA fragments to prepare them for dual indexing. A subsequent PCR amplification step was then used to add the index adapter sequences to create the final cDNA library. The adapter indices enabled the multiplexing of the libraries, which were pooled prior to cluster generation using a cBot instrument. The loaded flow-cell was then paired-end sequenced (76 + 76 cycles, plus indices) on an Illumina HiSeq4000 instrument. Finally, the output data was demultiplexed and BCL-to-Fastq conversion performed using Illumina’s bcl2fastq software, version 2.20.0.422 (Illumina, Inc., San Diego, CA, USA).

For RNAseq analysis, the unmapped paired-end sequences from the HiSeq 4000 were assessed by FastQC (http://www.bioinformatics.babraham.ac.uk/projects/fastqc/, accessed on 15 October 2021). Sequence adapters were removed, and reads were quality trimmed to quality q20 using Trimmomatic_0.36 (PMID: 24695404). The reads were mapped against the reference human (hg38) genome and counts per gene were calculated using annotation from GENCODE 30 (http://www.gencodegenes.org/, accessed on 15 October 2021) using STAR_2.5.3a (PMID: 23104886). Downstream normalisation, principal components analysis (PCA) and differential expression were calculated in DESeq2_1.20.0 using default settings (PMID:25516281). Differentially expressed genes (DEGs) were identified between NC-SMCs derived from CADASIL patients and controls (isoCtrl and wild-type) using an adjusted p < 0.05 and |log₂(fold change)| > 1.

Results were visualized using heatmaps generated with the pheatmap package (v1.0.12; R Foundation for Statistical Computing, Vienna, Austria). The volcano plots derived from DESeq2 statistical outputs was plotted by https://www.bioinformatics.com.cn (last accessed on 10 Dec 2024), an online platform for data analysis and visualization.Gene Ontology (GO) and Kyoto Encyclopedia of Genes and Genomes (KEGG) pathway enrichment analyses, as well as Gene Set Enrichment Analysis (GSEA), were conducted using the clusterProfiler package (v3.16.0; Bioconductor, Boston, MA, USA) to characterize biological processes and signalling pathways enriched among significantly upregulated or downregulated genes, and the pictures were plotted by https://www.bioinformatics.com.cn

### Statistical analysis

All statistical analyses were performed using GraphPad Prism version 10.0 (GraphPad Software, San Diego, CA, USA) and R version 4.4.2 (R Foundation for Statistical Computing, Vienna, Austria). Data are presented as mean ± standard error of the mean (SEM).

The Shapiro-Wilk test was employed to assess the normality of data distributions. For comparison between CADASIL patient and their respective isoCtrls, unpaired Student’s t-test was utilised. One-way ANOVA followed by Tukey’s post hoc test was applied for multiple comparison amongst the three lineage-specific VSMCs. For experiments involving two independent variables, *NOTCH3* variants and treatment, two-way ANOVA followed by Tukey’s multiple comparisons test was conducted to evaluate group differences. A p-value of less than 0.05 was considered statistically significant. All experiments were conducted with a minimum of three biological replicates as three independent rounds of differentiation of each iPSC line. Detailed sample sizes and the number of replicates for each experiment are specified in the corresponding method.

## Acknowledgement

The study is partially supported by the British Heart Foundation (PG/12/31/2952). We thank Dr Adam Dickinson for his contribution to the establishment of the iPSC lines. We thank Dr. Helen Murphy and Iris Trender-Gerhard from Manchester Centre for Genomic Medicine for help in recruiting and taking biopsies from CADASIL patients, and Meenakshi Minnis and Stephen Trueman of MCGM for help in growing HDFs and karyotyping iPSCs. We would like to thank Prof Richard Baines for his PhD supervising role and valuable suggestions. We would like to thank the University of Manchester Bioimaging Core Facility for guidance on confocal microscopy and sample preparation. We wish to thank Dr Aleksandr Mironov in the FBMH EM Core Facility (RRID:SCR_021147) for his assistance and the Wellcome Trust for equipment grant support to the EM Facility. The authors would like to thank Rachel Scholey and Andy Hayes of the Bioinformatics and Genomic Technologies Core Facilities at the University of Manchester for providing support regarding RNAseq. We thank Prof Susan J Kimber’s group for sharing their colony isolating facilities. We are grateful to Shi-yang Li for his assistance with the RNAseq analysis using R code. CY is supported by a Post-Doctoral Fellowship from the China Scholarship Council (ref No.201908500055).

## Author contributions

T.W. conceptualised and designed study, supervised the project, and manuscript writing. X.Z. performed most experiments and conducted data acquisition and analysis; contributed to manuscript writing. C.Y. and A.A: contributed to the generation of isogenic controls using CRISPR/Cas9. A.Z and H.Z: Developed AI tools for imaging analysis. P.S. Contributed to CADASIL patient recruitment and skin biopsy and manuscript proofreading.

