## Supplemental figures and table for "Selective vulnerability of cerebral vasculature to *NOTCH3* variants in small vessel disease and rescue by phosphodiesterase-5 inhibitor"

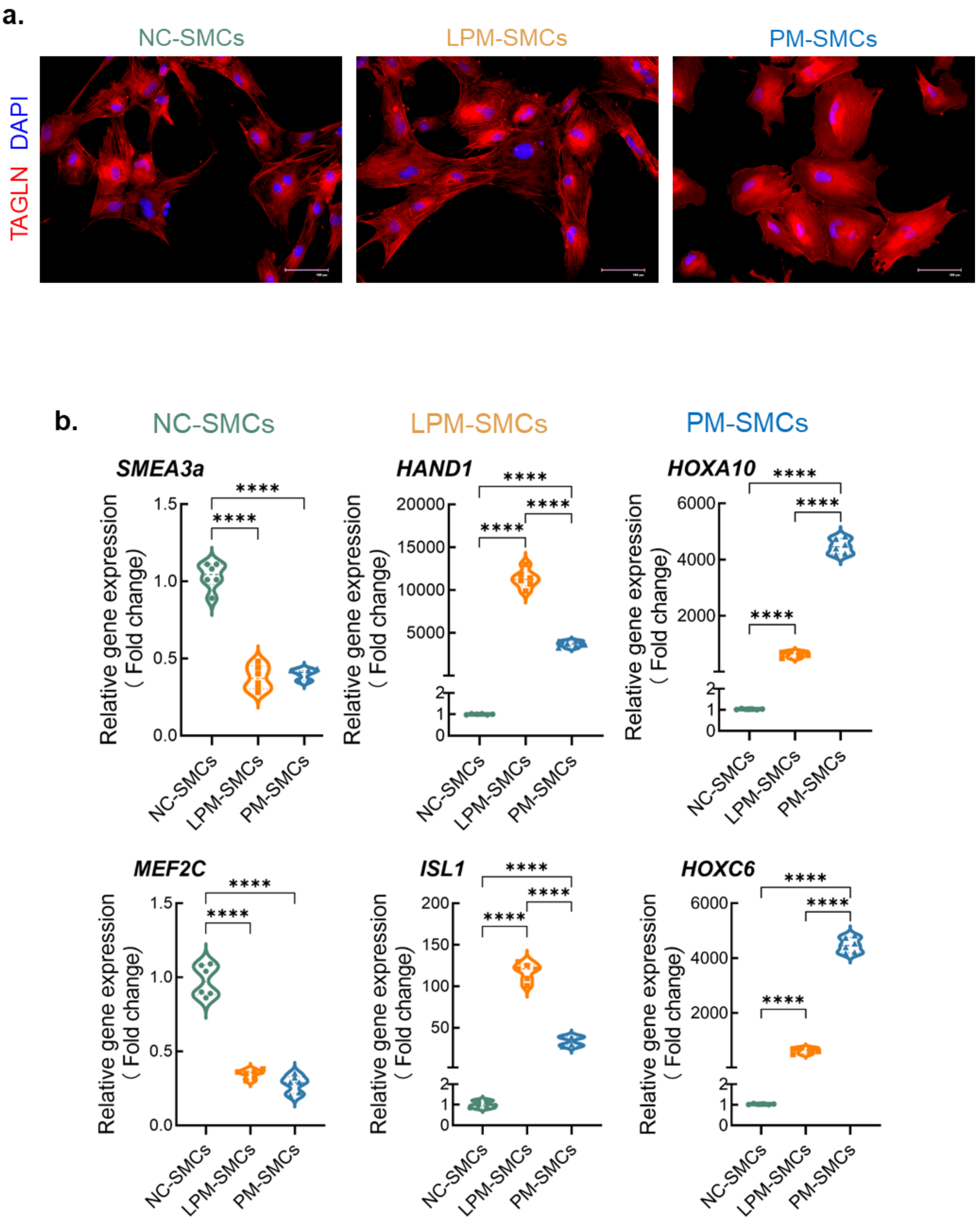


**Figure S1. The verification of iVSMCs.** IPSCs were differentiated into VSMCs via neural crest lineage (NC-SMCs), lateral plate mesoderm lineage (LPM-SMCs) and paraxial mesoderm lineage (PM-SMCs). **a** Lineage-specific VSMCs on differentiation day 18 were immunofluorescent stained for VSMC marker SM22α (red). Nuclei were counterstained by DAPI (blue). Scale bar = 100μm. **b** RT-qPCR results demonstrating that each type of the VSMCs expressed the corresponding lineage-specific markers: *SEMA3α* and *MEF3c* for NC-SMCs, *ISL1* and *HAND1* for LPM-SMCs, and *HOXA10* and *HOXC6* for PM-SMCs. Data are presented as mean ± SEM from 3 independent iPSC differentiations (n=3). One-way ANOVA and Tukey's post hoc test, ****p ≤ 0.0001.

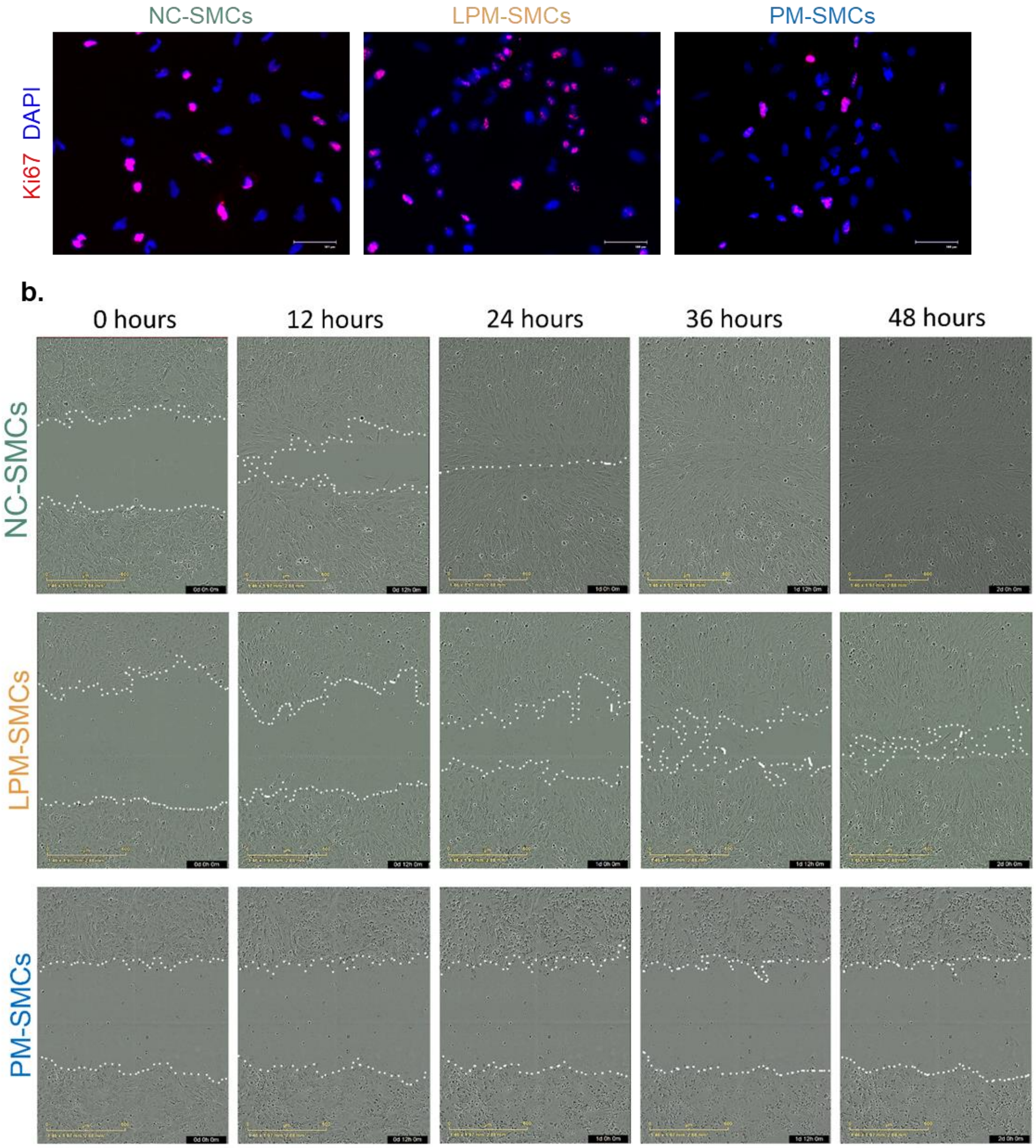


**Figure S2. Comparison of proliferation and migration between lineage-specific VSMCs derived from iPSCs.** IPSCs were differentiated into VSMCs via neural crest lineage (NC-SMCs), lateral plate mesoderm lineage (LPM-SMCs) and paraxial mesoderm lineage (PM-SMCs). **a** Cell proliferation was determined by ki67 immunofluorescent staining. Scale bar = 100 µm**. b** Cell migration was determined by IncuCyte live cell imaging of wound healing assay over 48 hours. Scale bar = 500 µm.

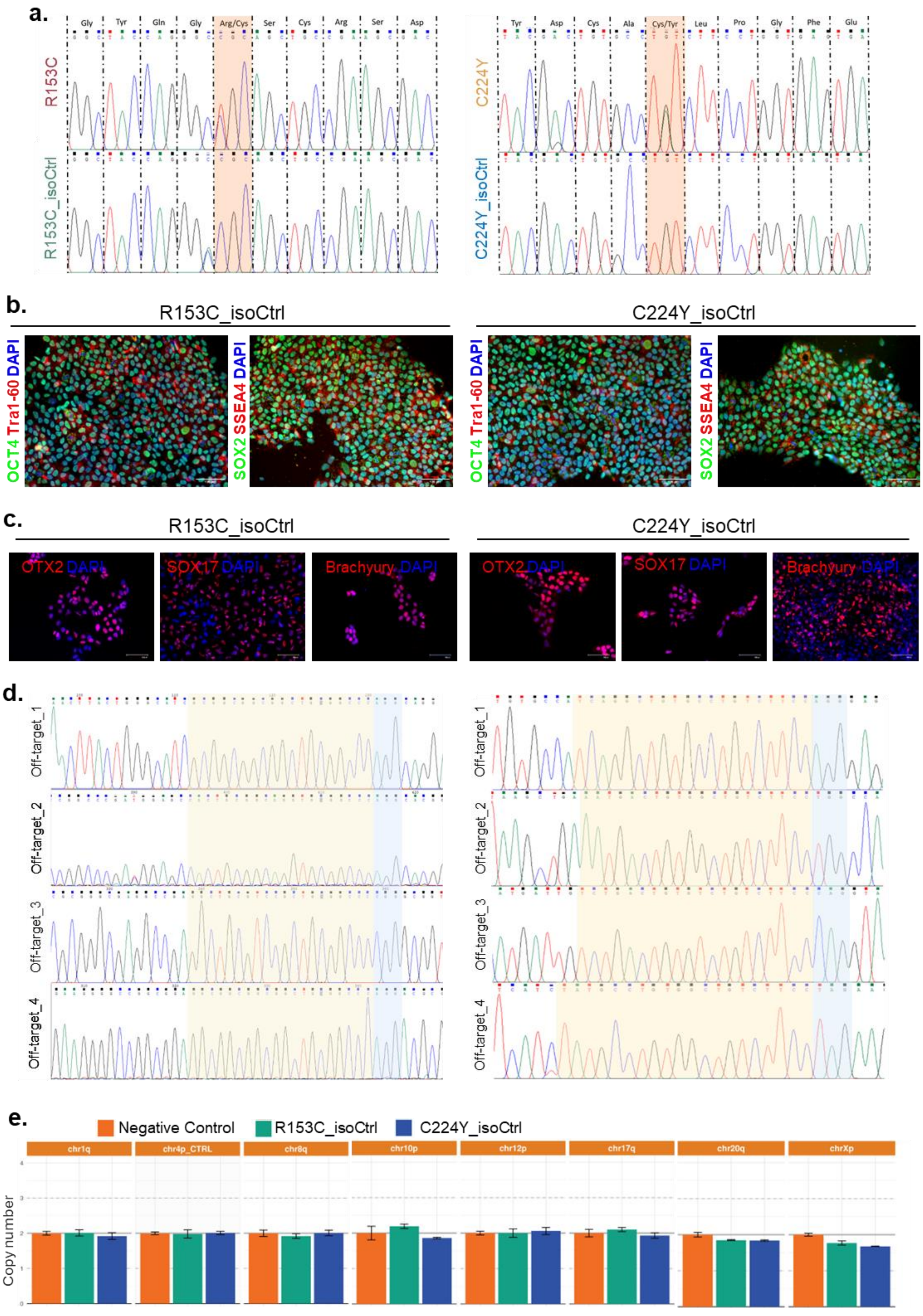

a.
Gly Tyr Gln Gly Arg/Cys Ser Cys Arg Ser Asp
R153C
R153C_isoCtrl
Tyr Asp Cys Ala Cys/Tyr Leu Pro Gly Phe Glu
C224Y
C224Y_isoCtrl
b.
R153C_isoCtrl
C224Y_isoCtrl
OCT4 Tra1-60 DAPI
SOX2 SSEA4 DAPI
c.
R153C_isoCtrl
C224Y_isoCtrl
OTX2 DAPI
SOX17 DAPI
Brachyury DAPI
d.
Off-target_1
Off-target_2
Off-target_3
Off-target_4
e.
Negative Control
R153C_isoCtrl
C224Y_isoCtrl
chr1q
chr4p_CTRL
chr8q
chr10p
chr12p
chr17q
chr20q
chrXp
Copy number

**Figure S3. Creation and characterisation of isogenic control iPSC lines.** IPSC lines with CADASIL *NOTCH3* variants R153C and C244Y, respectively, were subjected to CRISPR/Cas9 gene editing to generate corresponding isogenic control iPSC lines (isoCtrl) by correcting the two *NOTCH3* variants. **a** Chromatograph of DNA sequencing demonstrating successful changes of R153C and C224Y back to wild type. **b** Immunofluorescent staining of pluripotency markers OCT4 (green), SOX2 (green), Tra1-60 (red) and SSEA4(red). Nuclei were overstained by DAPI (blue). Scale bar = 100 µm. **c** Immunofluorescent staining of cells from trilineage differentiation of the isogenic control iPSC lines demonstrating the presence of ectodermal marker OTX2 (red), endodermal marker SOX17 (red), and mesodermal marker Brachyury (red), which confirm the pluripotency of the iPSC lines. Nuclei were overstained by DAPI (blue). Scale bar = 100 µm. **d** DNA sequencing of the top four potential CRISPR off-targets sites confirming lack of off-target DNA editing. **e** hPSC Genetic Analysis Kit (StemCell) that detects the 8 most common karyotypic abnormalities reported in human iPSCs in the indicated chromosomes (chr) showing normal copy numbers in each chromosome.

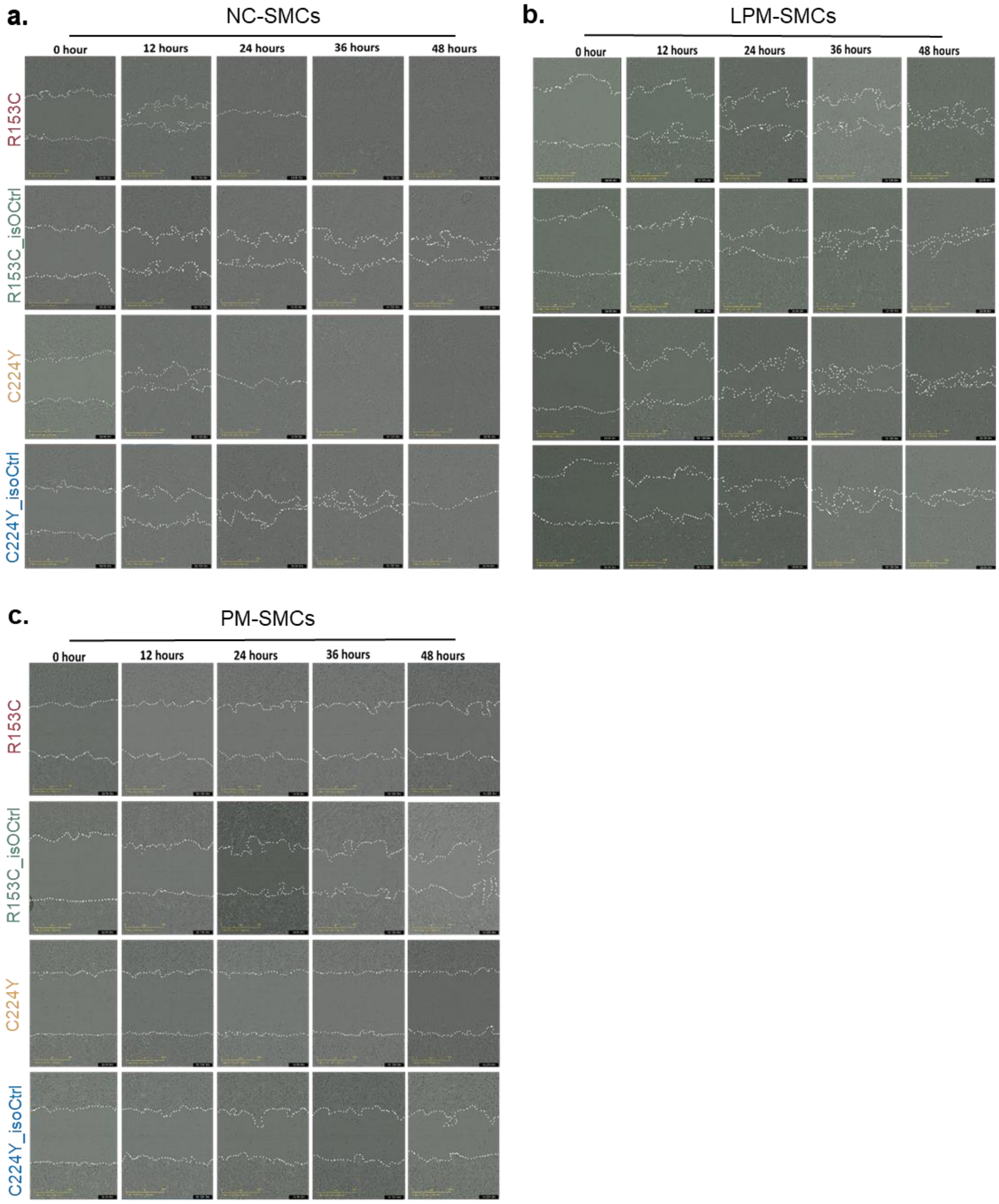


**Figure S4. Migration of iPSC derived VSMCs by wound healing assay.** IPSCs from two CADASIL patients (R153C and C224Y) and their isogenic controls (isoCtrl) were differentiated into VSMCs via neural crest lineage (NC-SMCs), lateral plate mesoderm lineage (LPM-SMCs) and paraxial mesoderm lineage (PM-SMCs). Cell migration was determined by IncuCyte live cell imaging of wound healing assay. Figures show light microscopic images of the migration over 48 hours. Scale bar = 500 μm.

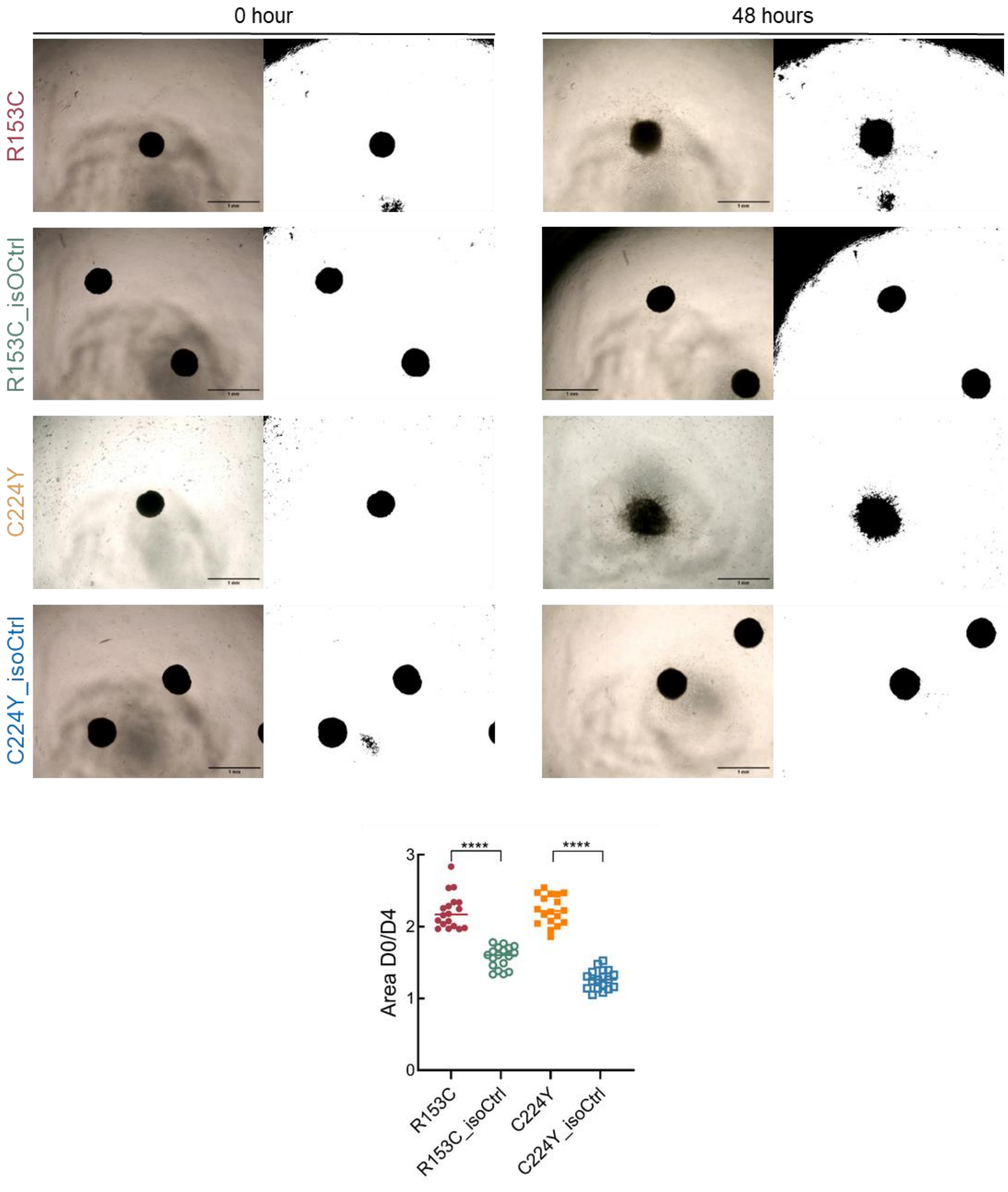


**Figure S5. Migration of iPSC derived VSMCs by spheroid migration assay.** IPSCs from two CADASIL patients (R153C and C224Y) and their isogenic controls (isoCtrl) were differentiated into iVSMCs via neural crest lineage and grew as spheroids to allow cells to migrate outwards for 48 hours. Results were quantified by measuring spheroid areas. Scale bar = 500 μm.

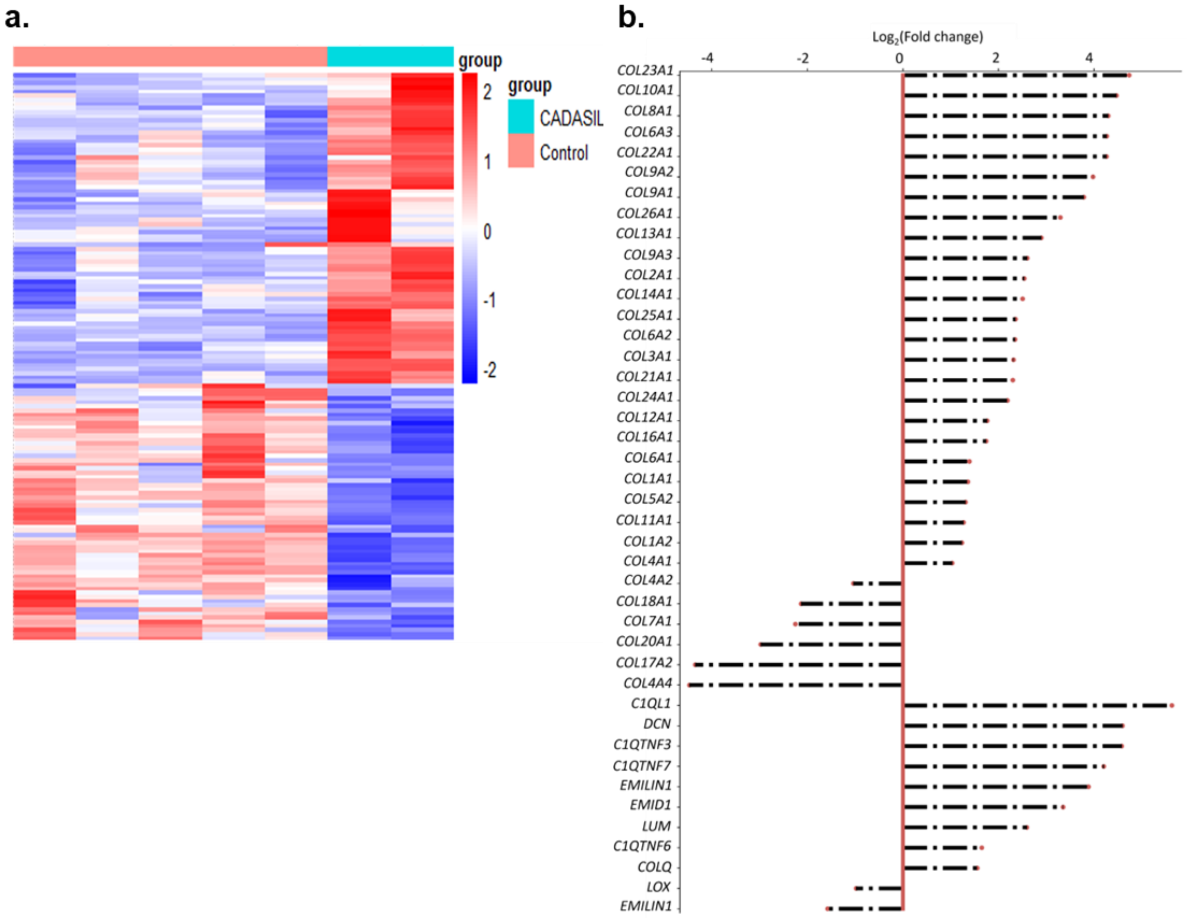


**Figure S6. Analysis of RNAs sequencing results on extracellular matrix (ECM) related genes.** IPSCs from two CADASIL patients (R135C & C224Y) and respective isogenic controls (isoCtrl) were differentiated into iVSMCs via neural crest lineage which were subjected to RNA sequencing. **a** Heatmap showing differentially expressed ECM genes between iVSMCs of the two patients and their isogenic controls under the gene ontology (GO) term "extracellular matrix organisation". **b** Differentially expressed genes related to collagen.

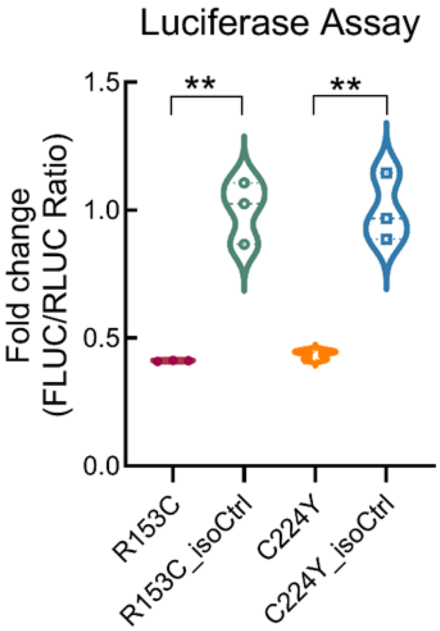


**Figure S7. Notch signalling luciferase assay.** IPSCs from two CADASIL patients (R153C and C224Y) and their isogenic controls (isoCtrl) were differentiated into iVSMCs via neural crest lineage and subjected to Dual-Luciferase reporter assay. Results are presented as fold changes of the ratio of Firefly luciferase activity to Renilla luciferase activity (FLUC/RLUC). Data are mean ± SEM from 3 independent iPSC differentiations (n=3). Unpaired Student's *t* test, **p ≤ 0.01.

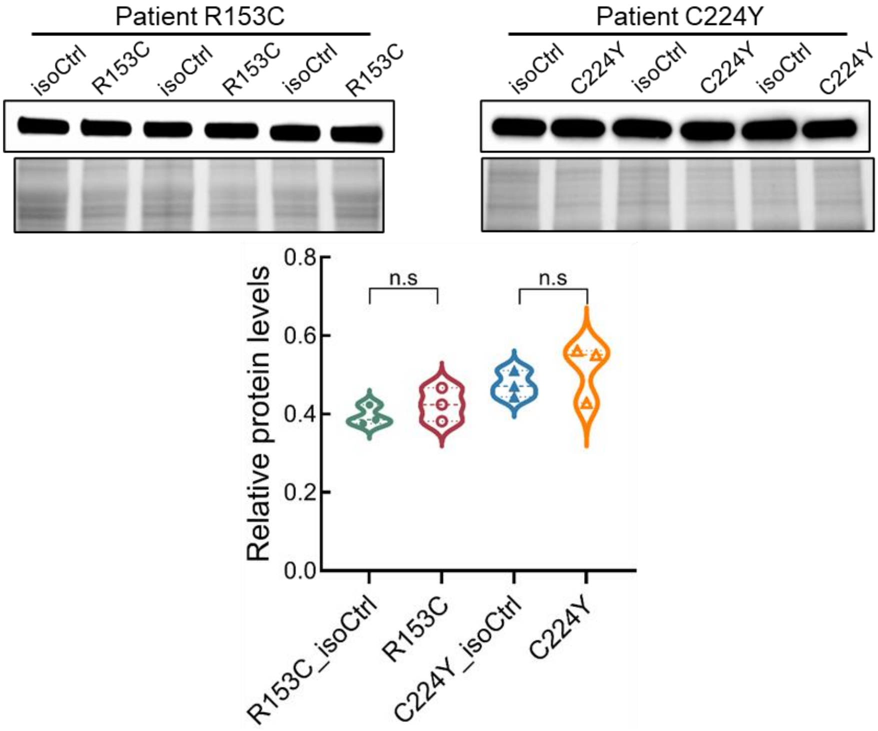


**Figure S8. Western blotting of inducible nitric oxide synthase (iNOS).** IPSCs from two CADASIL patients (R153C and C224Y) and their isogenic controls (isoCtrl) were differentiated into iVSMCs via neural crest (NC) lineage and subjected to western blotting to determine cellular iNOS protein levels. Results were quantified and normalised to the total protein loading control (representative areas are shown under each blot). under each blot. Data are mean ± SEM from 3 independent iPSC differentiations (n=3). Unpaired Student's *t* test, n.s., no significant difference.

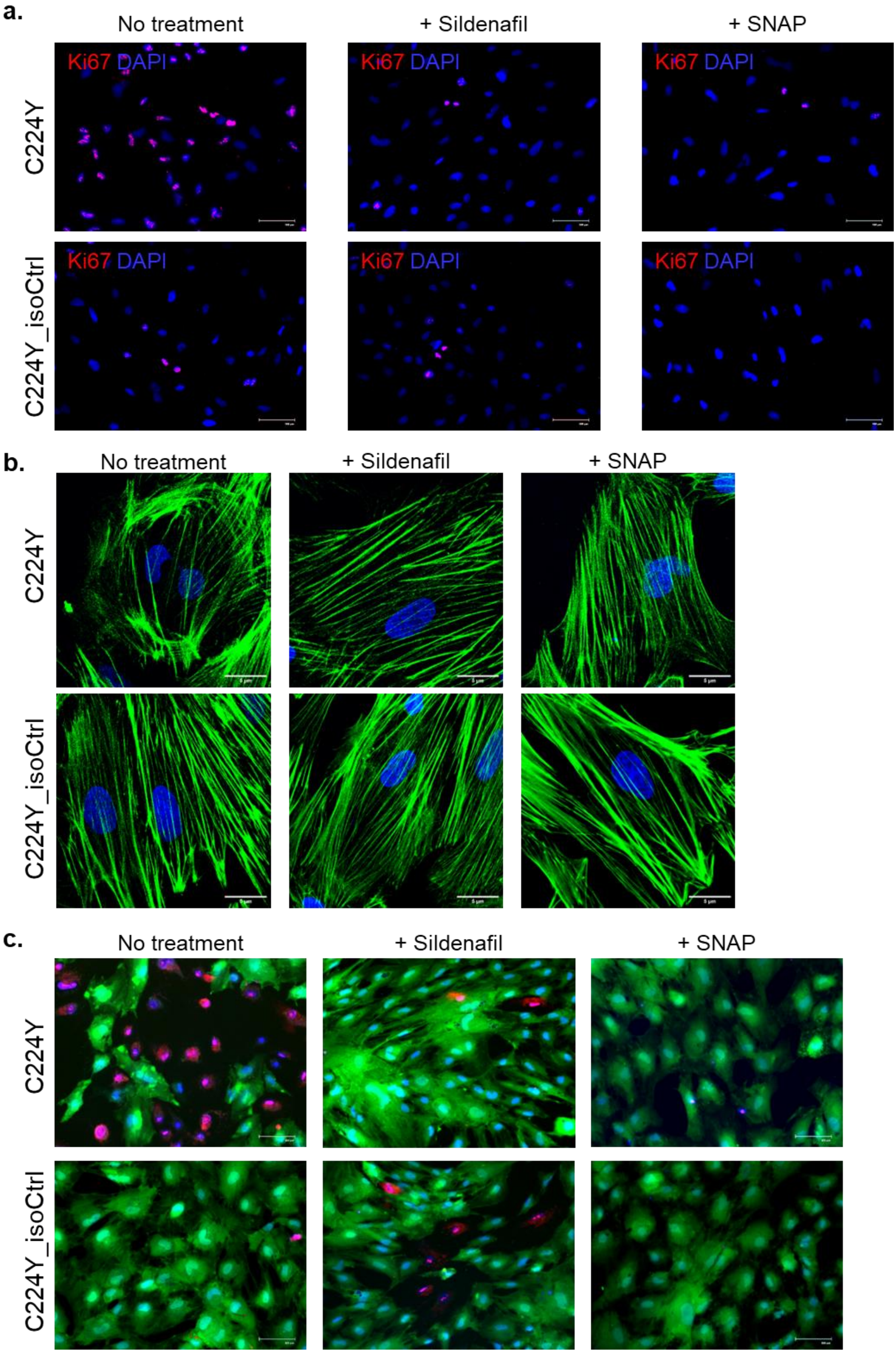
a.
No treatment
+ Sildenafil
+ SNAP
C224Y
C224Y_isoCtrl
Ki67 DAPI
b.
No treatment
+ Sildenafil
+ SNAP
C224Y
C224Y_isoCtrl
c.
No treatment
+ Sildenafil
+ SNAP
C224Y
C224Y_isoCtrl

**Figure S9. SNAP and Sildenafil rescue of abnormal proliferation, contraction and cell death of CADASIL iVSMCs.** IPSCs from two CADASIL patients (R135C & C224Y) and their isogenic controls (R153C_isoCtrl and C224Y_isoCtrl) were differentiated into iVSMCs via neural crest (NC) lineage. **a** The iVSMCs were subjected to proliferation assay by Ki67 immunofluorescent staining in the presence or absence of the nitric oxide (NO) donor (SNAP) or PDE5 inhibitor (Sildenafil). **b** F-actin staining of iVSMCs derived from iPSCs of CADASIL and isoCtrls treated with or without SNAP or Sildenafil. **c** Live/dead staining of iVSMCs derived from iPSCs of CADASIL and isoCtrls using Calcein-AM (green) and BOBO-3 iodide (red), respectively. Scale bars = 100 μm. **a** and **c**; 5 μm in **b.** Nuclei were counterstained by DAPI.

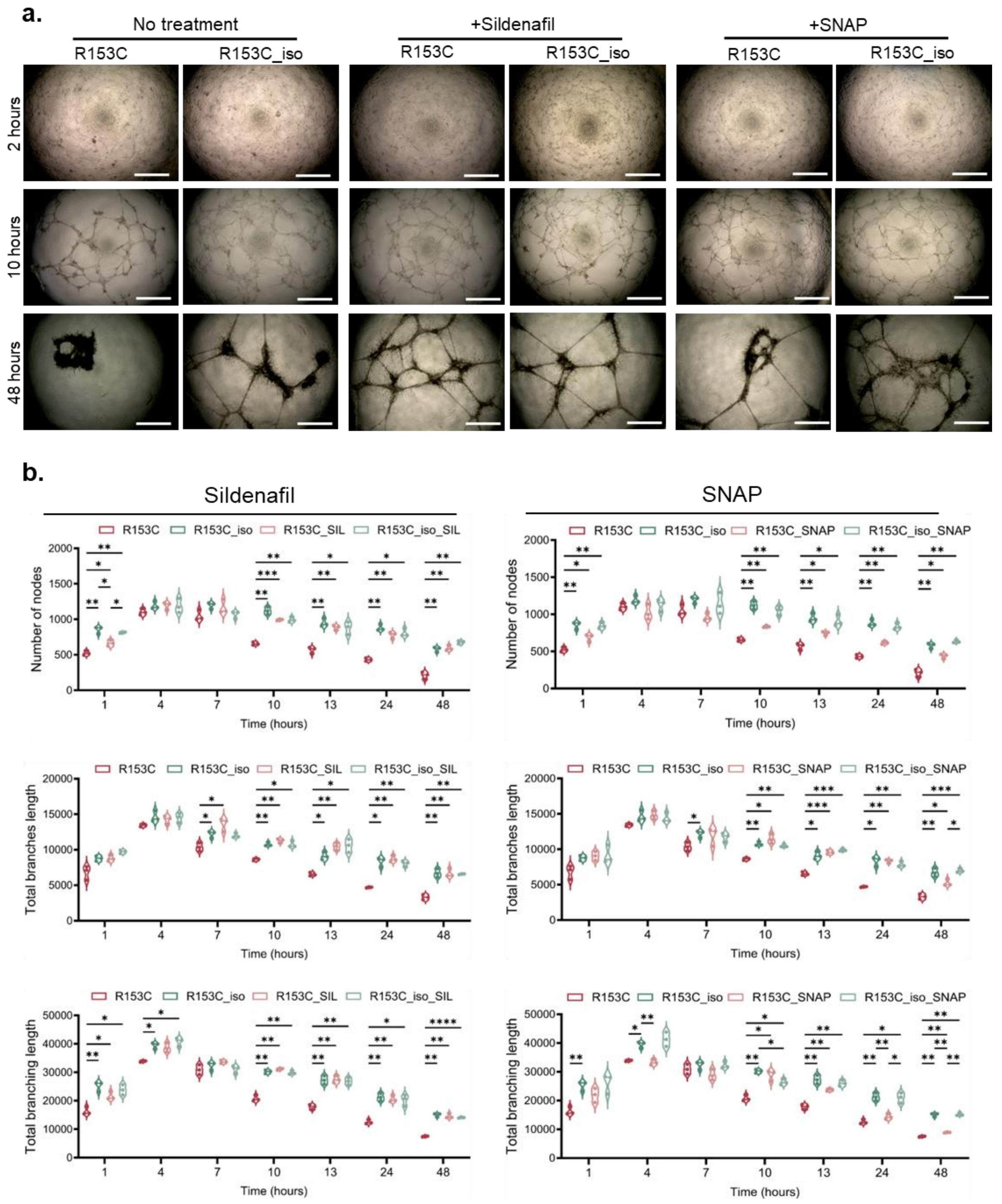


**Figure S10. SNAP and Sildenafil rescue of impaired *NOTCH3*-R153C iVSMCs in supporting angiogenesis.** IPSCs from CADASIL patient R135C and the isogenic control line (R153C_isoCtrl) were differentiated into iVSMCs via neural crest (NC) lineage. The iVSMCs were mixed with HUVECs in a 1:2 ratio and subjected to angiogenesis in Matrigel for 24 hours to determine the capability of iVSMCs in supporting the angiogenic network structure. **a** Images of the angiogenic network formation under light microscope in each condition. Scale bar = 1 mm. **b** Quantification of the angiogenesis results at each time point by measuring number of nodes, total branches length, and total branching length using ImageJ. Data are mean ± SEM from 3 independent iPSC differentiations (n=3). Two-way ANOVA and Tukey's post hoc test, *p ≤ 0.05, **p ≤ 0.01, ***p ≤ 0.001, and ****p ≤ 0.0001.

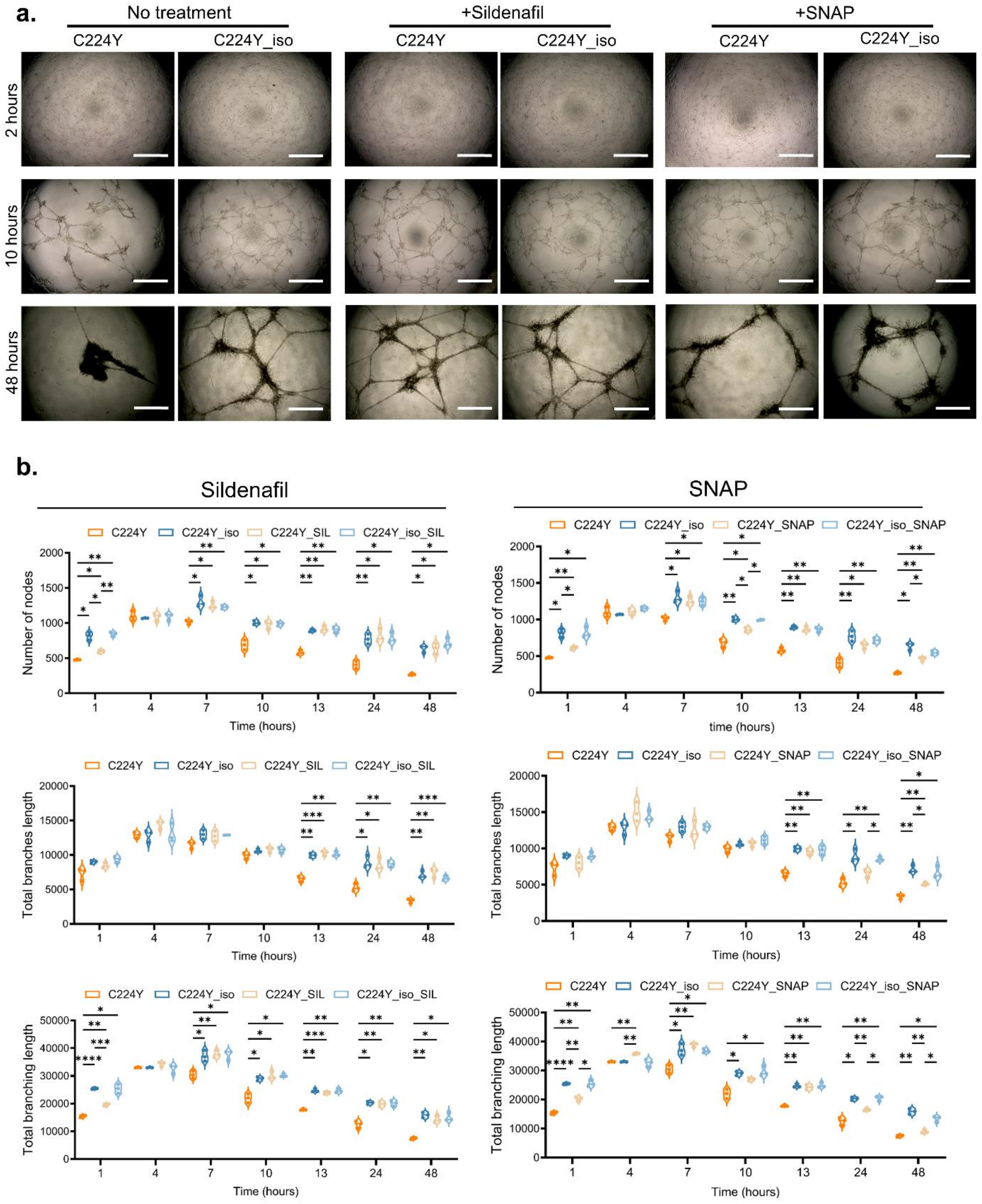


**Figure S11. SNAP and Sildenafil rescue of impaired *NOTCH3*-C224Y iVSMCs in supporting angiogenesis.** IPSCs from CADASIL patient C224Y and the isogenic control line (C224Y_isoCtrl) were differentiated into iVSMCs via neural crest (NC) lineage. The iVSMCs were mixed with HUVECs in a 1:2 ratio and subjected to angiogenesis in Matrigel for 24 hours to determine the capability of iVSMCs in supporting the angiogenic network structure. **a** Images of the angiogenic network formation under light microscope in each condition. Scale bar = 1 mm. **b** Quantification of the angiogenesis results at each time point by measuring number of nodes, total branches length, and total branching length using ImageJ. Data are mean ± SEM from 3 independent iPSC differentiations (n=3). Two-way ANOVA and Tukey's post hoc test, *p ≤ 0.05, **p ≤ 0.01, ***p ≤ 0.001, and ****p ≤ 0.0001.

### Table S1. PCR primers used in the study.

| Primer | Forward | Reverse | Efficiency |
|---|---|---|---|
| ***GAPDH*** | CATGTTCGTCATGGGTGTGAACCA | ATGGCATGGACTGTGGTCATGAGT | 94.92 |
| ***ACLP*** | ACCCACACTGGACTACAATGA | GTTGGGGATCACGTAACCATC | 96.06 |
| ***ACTA2*** | TGACAATGGCTCTGGGCTCTGTAA | TTCGTCACCCACGTAGCTGTCTTT | 100.8 |
| ***ACTG2*** | ATTGTGCGAGACATCAAGGAG | CCATGCCAATAAAGGAAGGCT | 95.26 |
| ***ACTN4*** | GCAGCATGGGCGACTACAT | TTGAGCCCGTCTCGGAAGT | 94.25 |
| ***CDH13*** | AGTGTTCCATATCAATCAGCCAG | CGAGACCTCATAGCGTAGCTT | 88.90 |
| ***CDH2*** | TGCGGTACAGTGTAACTGGG | GAAACCGGGCTATCTGCTCG | 100.54 |
| ***CNN1*** | GTCCACCCTCCTGGCTTT | AAACTTGTTGGTGCCCATCT | 99.21 |
| ***COL1A1*** | ATCAACCGGAGGAATTTCCGT | CACCAGGACGACCAGGTTTTC | 108.49 |
| ***COL1A2*** | GTTGCTGCTTGCAGTAACCTT | AGGGCCAAGTCCAACTCCTT | 87.31 |
| ***COL2A1*** | TGGACGCCATGAAGGTTTTCT | TGGGAGCCAGATTGTCATCTC | 105.8 |
| ***COL3A1*** | TTGAAGGAGGATGTTCCCATCT | ACAGACACATATTTGGCATGGTT | 97.20 |
| ***COL4A1*** | GGACTACCTGGAACAAAAGGG | GCCAAGTATCTCACCTGGATCA | 106.67 |
| ***COL4A2*** | TTATGCACTGCCTAAAGAGGAGC | CCCTTAACTCCGTAGAAACCAAG | 99.25 |
| ***FN*** | CGGTGGCTGTCAGTCAAAG | AAACCTCGGCTTCCTCCATAA | 98.19 |
| ***GUCY1A3*** | TCAGCCCTACTTGTTGTACTCC | CAGAATAGCGATGTGGGAATCAC | 89.27 |
| ***GUCY1B3*** | TGCTGGTGATCCGCAATTAC | CCAGGACACGCAAGATTGTATC | 96.28 |
| ***HAND1*** | CCATGCTCCACGAACCCTTC | CCTGGCGTCAGGACCATAG | 99.28 |
| ***HOX10A*** | CTCGCCCATAGACCTGTGG | GTTCTGCGCGAAAGAGCAC | 92.74 |
| ***HOX6C*** | ACAGACCTCAATCGCTCAGGA | AGGGGTAAATCTGGATACTGGC | 88.08 |
| ***ICAM*** | GTATGAACTGAGCAATGTGCAAG | GTTCCACCCGTTCTGGAGTC | 100.08 |
| ***ISL1*** | AGATTATATCAGGTTGTACGGGATC | ACACAGCGGAAACACTCGAT | 87.79 |
| ***ITGA1*** | GTGCTTATTGGTTCTCCGTTAGT | CACAAGCCAGAAATCCTCCAT | 89.91 |
| ***ITGA2*** | AGGTGGGGTTAATTCAGTATGCC | GATGTCTGGGATGTTGCTACAA | 95.03 |
| ***ITGA3*** | TCAACCTGGATACCCGATTCC | GCTCTGTCTGCCGATGGAG | 100.54 |
| ***ITGA4*** | CACAACACGCTGTTCGGCTA | CGATCCTGCATCTGTAAATCGC | 92.86 |
| ***ITGA5*** | AGACATTCGATCCCTCTACAACT | AATCGGCCAAACTCATCATGG | 103.00 |
| ***ITGA6*** | ATGCACGCGGATCGAGTTT | TTCCTGCTTCGTATTAACATGCT | 98.68 |
| ***ITGA7*** | CTGACTCCATGTTCGGGATCA | CACCTGTGAAGGTTTGGCG | 100.42 |
| ***ITGB1*** | CCTACTTCTGCACGATGTGATG | CCTTTGCTACGGTTGGTTACATT | 90.77 |
| ***ITGB3*** | CATGAAGGATGATCTGTGGAGC | AATCCGCAGGTTACTGGTGAG | 86.63 |
| ***MMP3*** | CGGTTCCGCCTGTCTCAAG | CGCCAAAAGTGCCTGTCTT | 83.65 |
| ***MMP9*** | TGTACCGCTATGGTTACACTCG | GGCAGGGACAGTTGCTTCT | 86.92 |
| ***MSN*** | GAGGATGTGTCCGAGGAATTG | GTCTCAGGCGGGCAGTAAA | 90.26 |
| ***MYH11*** | GACTTCCCTGCTCAATGCCT | GGACCTCTTCTCGTGGTTGG | 101.65 |
| ***MYH10*** | TGGTTTTGAGGCAGCTAGTATCA | AGTCCTGAATAGTAGCGATCCTT | 99.28 |
| ***MYL6*** | ACCAGACCGCAGAGTTCAAGGAG | CTCAAAGTCCAGCACCTTCACATT | 91.99 |
| ***MYL9*** | CCACATCCAATGTCTTCGCAATG | TGAAGTTGCCTTTCTTATCAATGGG | 89.74 |
| ***MYLK*** | GAGGTGCTTCAGAATGAGGACG | GCATCAGTGACACCTGGCAACT | 109.21 |
| ***MYOCD*** | CCACCTATGGACTCAGCCTAC | CTCAGTGGCGTTGAAGAAGAG | 100.20 |
| ***OCT4*** | AGACCATCTGCCGCTTTGAG | GCAAGGGCCGCAGCTT | 91.63 |
| ***PRKG1*** | CTTGGAGCTGTCGCAGATCC | TCTTTGATGATGCAACTGTCCTT | 97.21 |
| ***RBP1*** | CGCACGCTGAGCACTTTTAG | GCACTTGCGGTCATCTATGC | 105.26 |
| ***SEMA3a*** | GTGCCAAGGCTGAAATTATCCT | CCCACTTGCATTCATCTCTTCT | 90.04 |
| ***SMTN*** | CGGCTGCGCGTGTCTAATCC | CTGTGACCTCCAGCAGCTTCCG | 98.84 |
| ***SOX1*** | CCTCCGTCCATCCTCTG | AAAGCATCAAACAACCTCAAG | 91.29 |
| ***SPEG*** | AACCGCCGTTCTTCTGACAC | TGGTCCATAAGTGAGACCTTGAA | 97.26 |
| ***SPP1*** | CTCCATTGACTCGAACGACTC | CAGGTCTGCGAAACTTCTTAGAT | 93.63 |
| ***TAGLN*** | CGCGAAGTGCAGTCCAAAAT | CAGCTTGCTCAGAATCACGC | 85.01 |
| ***TBX6*** | CATCCACGAGAATTGTACCCG | AGCAATCCAGTTTAGGGGTGT | 100.11 |
| ***TIMP1*** | CCACCATGAGACCTCAACCC | GCCACTACAGCCGTATTCTCC | 92.65 |
| ***TIMP3*** | TGGCGCAGTGAGAACTTCG | CCCCGAGTAGAGGTCATCCAG | 97.32 |
| ***TFAP2A*** | AGGTCAATCTCCCTACACGAG | GGAGTAAGGATCTTGCGACTGG | 85.75 |
| ***VASP*** | ATGGCAACAAGCGATGGCT | CGATGGCACAGTTGATGACCA | 103.21 |
